# Epitome: Predicting epigenetic events in novel cell types with multi-cell deep ensemble learning

**DOI:** 10.1101/2021.06.10.447140

**Authors:** Alyssa Kramer Morrow, John Weston Hughes, Jahnavi Singh, Anthony Douglas Joseph, Nir Yosef

**Affiliations:** Electrical Engineering and Computer Science Department, University of California-Berkeley, 465 Soda Hall, Berkeley CA 94720-1776, United States; Computer Science Department, Stanford University, 353 Serra Mall, Stanford, CA 94305, United States; Center for Computational Biology, University of California-Berkeley, 108 Stanley Hall, Berkeley CA 94720-3220, United States; Unite Genomics, Inc., 1301 Marina Village Pkwy, Suite 320, Alameda CA 94501, United States

**Author notes:** Now at Google (Jahnavi Singh).

## Abstract

The accumulation of large epigenomics data consortiums provides us with the opportunity to extrapolate existing knowledge to new cell types and conditions. We propose Epitome, a deep neural network that learns similarities of chromatin accessibility between well characterized reference cell types and a query cellular context, and copies over signal of transcription factor binding and modification of histones from reference cell types when chromatin profiles are similar to the query. Epitome achieves state-of-the-art accuracy when predicting transcription factor binding sites on novel cellular contexts, and can further improve predictions as more epigenetic signals are collected from both reference cell types and the query cellular context of interest.

## 1 Introduction

Over the past decade, large scale projects such as ENCODE [1, 2] and the Roadmap Epigenomics project [3] have amassed over 20 terabytes of epigenetic profiles. These profiles include measurements of histone modifications, transcription factor binding, and chromatin accessibility, and are measured in a select set of cellular contexts. This data has enhanced our understanding of transcriptional regulation within these samples and has served to explore general questions in chromatin biology, pertaining to the function of the non-coding genome [4, 5], the interplay between histone modifications and transcription factors [6], and their association with transcription [7, 8, 9].

To date, one of the most prevalent types of genome-scale epigenetic information is chromatin accessibility. Respective assays such as DNase I hypersensitive sites sequencing (DNase-seq) [10] and Assay for Transposase-Accessible Chromatin using sequencing (ATAC-seq) [11] identify accessible chromatin regions genome-wide through enzymatic cleavage of exposed DNA. These assays do not only inform us about the local structure of chromatin, but also aid in identifying the location of genomic regions (mostly non-coding) that help regulate transcription from nearby loci [12, 13]. Beyond accessibility, regulatory regions are often associated with other epigenetic events such as binding of transcription factors (TFs) and other DNA binding proteins, or chemical modifications of histone tails [14]. These events are often indicative of the function of their respective region [2] and could be evaluated genome-wide through assays such as Chromatin Immunoprecipitation followed by DNA sequencing (ChIP-seq) [15] or Cleavage Under Targets and Release Using Nuclease (CUT&RUN)[16]. Although these assays can in principle be used to identify and characterize the activity of regulatory regions in a cellular context of interest [15], running separate ChIP-seq experiments for each of a large number of relevant DNA binding proteins and histone modifications is time and cost-intensive and, in some instances, unfeasible due to low input size.

As a result, the task of predicting the location of such epigenetic events in silico in lieu of experimental evaluation received great deal of attention [17]. Methods developed to predict such events can be broadly categorized based on the genomic properties they use as features for drawing predictions. The first broadly utilized category of classification methods is restricted to using only DNA sequences as features [18, 19, 20, 21]. As it has been repeatedly observed that the presence of a TF binding site may depend on a wider context around the site [22, 13], many approaches use neural networks to consider wide genomic contexts around a region of interest to predict epigenetic signal. Although these methods cannot predict epigenetic events that are specific to cellular contexts not seen during training, they can be used to explain the effect of changes in DNA sequence on the strength of the signal of DNA binding proteins and histone modifications. A second group of classification methods utilizes available measurements of chromatin state from a cellular context of interest to predict epigenetic events which were not measured experimentally [23, 24, 25, 26]. Many of these methods use chromatin accessibility in particular as an indication of cell type specificity due to its prevalence and ability to capture nuanced variation in accessibility at the protein-DNA physical interface [17, 27]. However, a subset of these methods augment the feature space to additionally incorporate DNA sequence information, RNA-sequencing information, or other epigenetic information specific to the cellular context of interest [24]. Regardless of the choice of features used, an important advantage of using epigenetic data as features is that it provides the means to distinguish epigenetic events between different cellular contexts that have a similar genome.

One common property of many classification methods designed to predict epigenetic signal in a new cellular context is that they learn a single model that is applied similarly to all positions in the genome [23, 24, 25, 26]. In this position-agnostic approach, models are applied separately for each candidate locus, using its local properties as features (such as the occurrence of DNA binding motifs or the enzymatic cleavage patterns of DNase-seq). To ensure accuracy and generalizability, these models are trained to identify local properties that are commonly predictive in many different loci. The natural caveat in this approach is that different loci may largely differ in terms of which specific features (or combination thereof) are in fact predictive. For instance, a single TF can bind the genome while interacting with different factors (i.e. by co-binding or tethered binding [28]), thus leading to different footprints and, possibly, different DNA binding motifs [29, 28].

In order to solve this caveat of position-agnostic learning, a third group of methods, referred to as imputation methods, directly uses epigenetic signal from known cellular contexts to predict in a new cellular context, instead of constructing complex features [30, 31, 32]. Novel imputation methods, such as Avocado [30] and PREDICTD [32], specifically use tensor factorization, assuming a low rank representation of the feature space, and jointly learn a model for all missing epigenetic signals [30, 32]. However, a caveat of joint learning of epigenetic signal in a single model is that objectives for each epigenetic signal can contradict each other, and update model parameters at different rates [33]. These limitations of joint learning can produce sub optimal predictions for epigenetic signal, compared to models optimized for a particular epigenetic signal of interest.

With these caveats in mind, we present **Epitome** - a conceptually simple alternative for predicting epigenetic events, such as TF binding and histone modifications. Similar to imputation methods, Epitome uses known epigenetic signal from multiple known cell types to predict epigenetic signal in a held out cellular context, bypassing the need to learn complex rules based on chromatin footprinting and DNA sequence. However, Epitome differs from imputation methods in three ways. First, Epitome approaches the problem of predicting epigenetic events as a classification task, treating each epigenetic signal as a binary event. As we explain in Section 4.3, this approach helps mitigate noise that arises from variation across different experiments, antibodies, and protocols that may affect quantitative results. Secondly, imputation methods such as Avocado [30] and PREDICTD [32] use a tensor factorization scheme, which assumes that epigenetic signals can be largely explained through a low dimensional representation, and looks for a single decomposition scheme for the entire tensor that couples all prediction tasks. Epitome has the flexibility of treating each prediction task completely independently from each other. As we show in Supplementary Figure S6(b), Epitome performs better when considering each epigenetic signal independently. Lastly, Epitome explicitly computes local similarities between the reference cell types and the query cellular context, and uses these as features in the model. These local similarities, along with information from reference cell types, are combined in an ensemble of reference cell types. This attribute helps the model explicitly learn the importance of experimental data from each reference cell type when predicting in the query cell type.

As input, Epitome requires chromatin accessibility in a query cellular context and a set of reference cell types in which the epigenetic event of interest was assayed. Epitome particularly requires chromatin accessibility as its primary indicator of cell type specificity because it is normally easy to generate [11, 10] and is informative of epigenetic events [34]. It then ”copies over” epigenetic events from the reference cell types to the query cellular context in positions where their chromatin accessibility is similar. In this case, we define similarity by comparing chromatin accessibility profiles in regions of different resolution surrounding the genomic locus in question. This similarity, along with the manner by which evidence across multiple reference cell types is aggregated, are learned using a neural network.

At the heart of our approach is the reliance on large amounts of publicly available data. While it is the case that practically any given cellular context will have a uniquely characteristic epigenome, there is a great deal of overlap between contexts. Consequently, epigenetic events that are uniquely observed in one cellular context become less prevalent as the number of other cellular contexts that have measured epigenetic events become available. This phenomenon is demonstrated in Figure 1(a), showing that for a given epigenetic event (binding of a certain TF, or a certain histone modification) the prevalence of genomic sites that are uniquely observed in only one cellular context in the ChIP-Atlas database [35] decreases substantially with the number of cellular contexts in which this event has been assayed with ChIP-seq. Particularly, we observe a mean level of coverage, or sensitivity, of over 90% for widely-assayed TFs and histone modifications (measured in 26 or more cellular contexts available in ChIP-Atlas). Looking ahead, we expect a continued increase in the size and quality of data sets in the public domain, leading to further increase in coverage.

**Figure 1:**
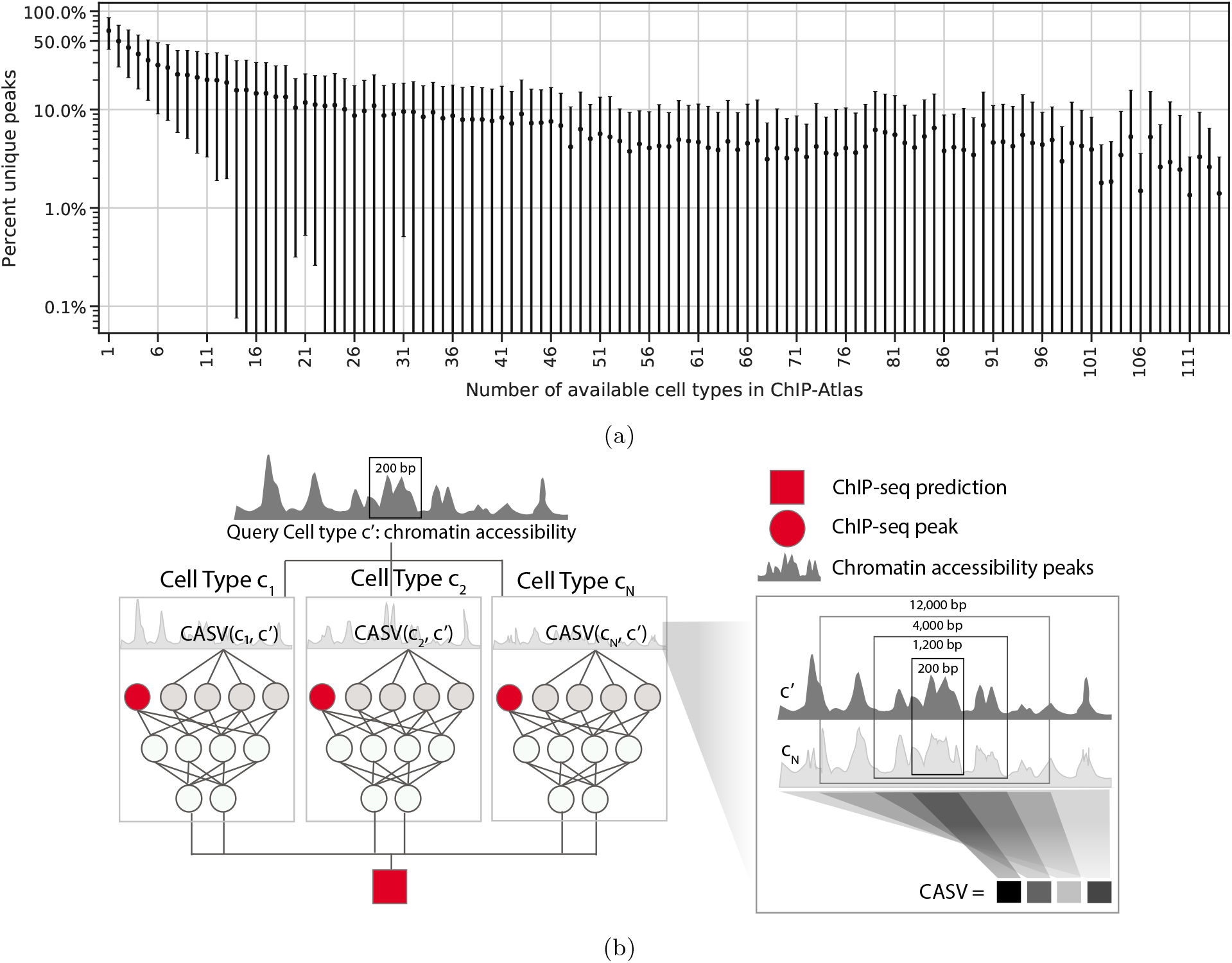
Epitome leverages multiple reference cell types in public data consortiums to learn the shared similarities of chromatin accessibility and its predictive effect on epigenetic events. (a) Weighted means and standard deviation of percent of unique peaks observed in a cell type as the number of available cell types for a given ChIP-seq target increases. Means and standard deviations are weighted inversely proportional to the number of data points for a given ChIP-seq target. ChIP-seq targets include transcription factors, histone modifications, chromatin accessibility, chromatin modifiers, and histones from called peaks in the ChIP-Atlas database [35]. (b) Schematic of Epitome for a single ChIP-seq target. Features for each cell type include ChIP-seq peaks at a genomic locus and the chromatin accessibility similarity vector (CASV), which compares the chromatin accessibility of each reference cell type to the query cellular context. The model outputs ChIP-seq peak probabilities for the query cellular context.

An important caveat of Epitome is that it will miss all sites that were not observed in any other cellular context. In the following, we show that in practical application this is a reasonable compromise. We compare Epitome to various methods that are designed to predict histone modifications and protein binding sites in novel cellular contexts. These methods include TF footprinting methods [23] and methods that use a combination of features constructed from chromatin accessibility, epigenetic signal, and DNA sequence [24, 36, 30]. Regardless of the choice of features used, Epitome achieves state-of-the-art accuracy when predicting TF binding sites in held out cellular contexts and chromosomes. We additionally demonstrate the deleterious effect of joint learning for multiple epigenetic signals and suggest that Epitome, along with current imputation methods, may achieve optimal performance by using a loss function dedicated to one epigenetic signal of interest. We additionally show how Epitome can extend its definition of cell type similarity to incorporate commonly assayed histone modifications, in additional to chromatin accessibility. We show that this extension of similarity between reference cell types and the query cellular context can further improve predictive performance.

Finally, we leverage Epitome to predict acetylation of H3K27 across seven time points in neural differentiation, demonstrating the ability of Epitome to leverage changes in chromatin accessibility from the same starting population to provide sensitive predictions of histone modifications over alternative methods. These results demonstrate that Epitome can significantly enrich the information that is routinely extracted in studies that use a limited view of the epigenome. Since such studies are increasingly prevalent in literature, we believe that such a method will have a substantial impact, providing a mechanistic complement to commonly used gene-ontology based approaches [37].

Epitome is an open-source project, and includes published tools and documentation. Epitome is available on GitHub and can be installed using the Python Package Index (PyPI). Documentation for Epitome is available at readthedocs.

## 2 Results

### 2.1 Epitome achieves state-of-the-art accuracy for prediction of TFBS

To evaluate Epitome, we compared to four state-of-the-art methods that predict transcription factor binding sites, where each method is a best-in-class representative of methods designed to predict TFBS. For each TF and each method, we evaluated the ability to predict the binding landscape in a held-out cellular context given information from other cellular contexts that were used for training. The first benchmark method, DeFCoM [23] represents the class of footprinting methods, which use enzymatic cleavage patterns of DNase 1 or Tn5 transposase as features to predict TFBS. Because DeFCoM and other footprinting methods are motif centric, they are only designed to predict binding in regions centered around motifs. We therefore compared Epitome and DeFCoM by evaluating the ability of each method to predict TFBS overlapping motifs for 77 TFs and chromatin modifiers across 40 cell lines, primary cells, and tissues (Supplementary Table S6). Figure 2(b) shows scatter plots of performance for both Epitome and DeFCoM in regions overlapping motifs, where we compare the area under the precision recall curve (auPRC) and partial area under the receiver operating characteristic curve (pAUC) (5% FPR). We note that while the former measure accounts for all candidate binding sites, the latter measure is meant to highlight the accuracy of top predictions [38]. On average, both methods perform comparably under both metrics, even though Epitome is not isolated to training in motif regions. Additionally, out of the 280 comparisons, Epitome ranks number one for 202 and 199 experiments for auPRC and pAUC, respectively, while DeFCoM ranks number one for only 78 and 76 experiments for auPRC and pAUC, respectively (there were 5 ties for pAUC between methods). These results suggest that Epitome can perform better than motif centric footprinting methods that use explicit knowledge of both motif location and cleavage patterns to predict TFBS.

**Figure 2:**
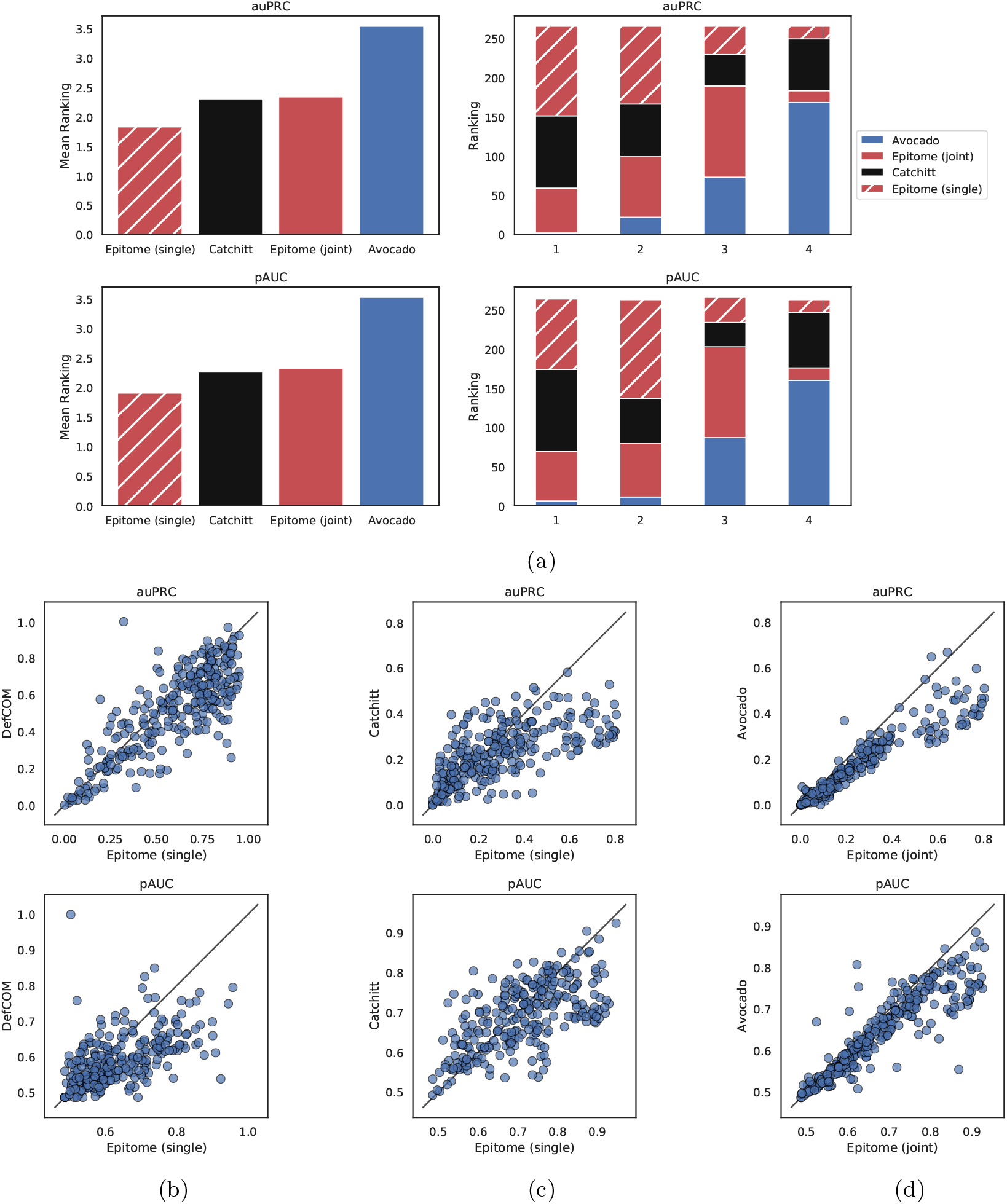
Comparison of methods for predicting transcription factor binding sites (TFBS) for 77 transcription factors (TFs) and chromatin modifiers in 40 primary cells, cell lines, and tissues from ENCODE. Transcription factor binding sites were predicted on all 200bp regions on chromosomes 8 and 9 that overlap at least one binding site in at least one of the 40 cell types considered. (a) Frequency at which each method obtains a rank for predicting TFBS across 77 transcription factors and chromatin modifiers in 40 held out cell lines, tissues, and primary cells, totaling 264 comparisons. Evaluated methods include Avocado [30], Catchitt [24], a joint Epitome model, and single Epitome models, where each TF is trained separately. (Left) Mean pAUC (5% FPR) and auPRC ranking for each method. (Right) Frequency at which each method obtains a rank based on pAUC and auPRC. (b) Scatter plots comparing auPRC and pAUC (5% FPR) between Epitome and DeFCoM. Only regions overlapping motifs specific to the TF being evaluated were considered. Both DeFCoM and Epitome trained individual models for each TF evaluated. (c) Scatter plots comparing auPRC and pAUC (5% FPR) between Epitome and Catchitt. Both Catchitt and Epitome trained individual models for each TF evaluated. (d) Scatter plots comparing auPRC and pAUC (5% FPR) between Epitome and Avocado. Both Avocado and Epitome trained joint models for all TFs evaluated.

Although many footprinting methods are constrained to only predict in regions overlapping motifs, TFs can bind in regions not overlapping canonical motifs associated with the TF of interest [29]. This is demonstrated in Supplementary Figure S5, which shows that the ratio of ChIP-seq peaks that do not overlap any motif for a TF in question can range from 0.45 to 1.0 for the 77 TFs and chromatin modifiers we evaluated (See Methods 4.5). Because limiting the prediction regime to sites that overlap motifs can eliminate many TFBS, Epitome and other methods have been designed to predict epigenetic events genome wide to eliminate this constraint. One example of such methods is Catchitt [24], a co-winner of the ENCODE-DREAM TFBS prediction challenge [36], which uses a combination of features including DNA sequence content, motif hits, expression levels of transcription factors (from RNA-seq), and chromatin accessibility (DNase-seq) to predict TFBS. Similar to DeFCoM, Catchitt considers accessibility data in a quantitative form, albeit at a default binned resolution of 50bp. It also uses a wider genomic context, taking up to 2,000 bp around the candidate site of interest. Additionally, Avocado [30] represents a class of imputation methods, designed to impute missing signal for cell types and assays that have not been measured experimentally. To compare to these methods, we first evaluated chromosome wide predictions of Epitome, Avocado, and Catchitt, on all 77 TFs and chromatin modifiers across 40 cell lines, primary cells, and tissues (Supplementary Figure S6) on held out chromosomes 8 and 9 in Figure 2(a), for a total of 264 comparisons. Here, we consider two versions of Epitome: single, which trains an individual model for each TF, and joint, which trains a single model to predict all TFs. These models are separately considered because Catchitt uses a single prediction approach, while Avocado uses a joint approach. Figure 2(a) shows that of the four methods, Epitome models trained individually for each TF perform the best for a majority of the 264 experiments. While joint Epitome models perform similarly in terms of pAUC and better for many instances in terms of auPRC compared to single Catchitt models (Supplementary Figure S6(a)), Avocado performs poorest on these metrics (Figure 2(a)). Supplementary Figure S6(b) additionally shows that for many cases, single Epitome models perform better than jointly trained Epitome models. It is unsurprising to note that single models often perform better than joint models. In these cases, learning objectives for different TFs can have complex or competing dynamics [33]. As shown in Supplementary Figure S16, different ChIP-seq targets trained in a joint model converge at different iterations during training. This difference in convergence across targets can result in variance in the level of fit for each ChIP-seq target. Regardless, we find that overall performance of both joint and single Epitome models is higher than the other two methods in the majority of the 264 experiments, based on both auPRC and pAUC metrics (Figure 2(a)).

We also consider a different group of methods that use NN architectures to predict TFBS from DNA sequence, but are not able predict on new cell types. This includes DeepSEA [19], which uses a convolutional neural network to learn the mapping from DNA sequence to TFBS. Surprisingly, we find that the accuracy of Epitome compares similarly or better to these types of models, which use all the data during training (i.e., not leaving out one of the cell line as query). This class of methods use DNA sequence alone as features to predict binding of transcription factors. While the goal of DeepSEA is to predict the effect of sequence mutations on the epigenetic events it is trained with (rather than predicting events in unobserved cellular contexts), the fact that it sees all data during training provides a conceptual upper bound for accuracy. To compare to DNA sequence based methods, we compared the performance of a set of 17 TFs which were assayed by the ENCODE consortium in four cell lines that were available in DeepSEA models, resulting in 68 comparisons. All data used is from ENCODE, and was aligned to the hg19 genome. Figure S10 shows that even though DeepSEA trains using a given test cell line and Epitome does not, Epitome performs comparably to DeepSEA in both pAUC and auPRC. These results demonstrate that Epitome can effectively generalize to unseen cellular contexts. Supplementary Figure S11 show receiver operating characteristic curves and precision-recall curves for these methods in all four held out cell lines from ENCODE.

### 2.2 Epitome places an upper bound on maximum achievable sensitivity

Because Epitome constructs features using binarized epigenetic events in reference cell types, models are a-priori limited to predict epigenetic events in regions in which the epigenetic event in question has been observed in a previous experiment. Although this limitation could be alleviated by using genome-wide continuous signal, which is available in all genomic regions, we explain in Section 4.3 our explicit choice to use binary epigenetic signal to reduce noise. In Figure 1(a), we calculated the fraction of unique peaks observed a held out cell type for all ChIP-seq targets available in ChIP-Atlas [35] to determine an upper bound for the sensitivity that could be achieved using Epitome, given this limitation. When more than 26 cell types are available for a given ChIP-seq target, we observe that sensitivity exceeds 90%, on average. Furthermore, we observe that the upper bound on sensitivity increases quickly as the number of available reference cell types increases (see Methods 4.2). While this approach places a strict upper bound on the achievable sensitivity, in Figure 2 we demonstrate that this strategy leads to a better overall balance between sensitivity and specificity, compared to existing methods.

### 2.3 Models trained on multiple cell types similar to the query cell type provide improved accuracy

Of the methods compared to in Section 2.1 that provide predictions of TFBS specific to a cellular context of interest, none are designed to jointly train on multiple reference data sets. This limitation bypasses the opportunity to jointly learn from multiple cell types and can generate misguided predictions when the query context greatly differs from the cell types selected for training. As previously mentioned, Figure 1(a) shows that as the number of cell types available for a given epigenetic event measured with ChIP-seq increases, the fraction of unique TF binding sites or histone modifications observed in a new cell type decreases. From this trend, we conclude that as more cell types for a given event are used to train a model, it is more likely that the model will have seen that event in the genomic location we are trying to predict. We therefore sought to understand the effect of the quantity and choice of reference cell types used for training Epitome on accuracy when predicting epigenetic signal in a new cellular context.

To assess the effect of the number of cell types used during training on model performance, we trained models for 82 different ChIP-seq targets, which included TFs, histones, and histone modifications. For each ChIP-seq target, we considered a range of models trained on different numbers of ENCODE cell types as references (ranging from 2 to *n* − 1 reference cell types, where *n* is the number of cell types available for the respective ChIP-seq target). Cell types used include cell lines, in-vitro differentiated cells, primary cells, and tissues from ENCODE [1]. For each ChIP-seq target and choice of cell type count used for training, we trained four models with different combinations of training and validation cell types. Each of these models were evaluated by predicting peaks across validation chromosome 7, which was held out from training. This procedure resulted in training 4 * (*n* − 2) models for each of the 82 ChIP-seq targets, where each model trained on a different number and combination of reference cell types.

Figures 3(a) and 3(b) demonstrate the change in performance (auPRC) in predicting TFBS with increasing numbers of reference data sets. Across the 59 TFs included in this analysis, we observe an overall consistent increase in performance as one considers larger numbers of reference cell types in which the TF in question was measured. While for most TFs, the number of cell types for which information was available is limited (under thirteen), the ENCODE collection includes over forty contexts for the CCCTC-Binding Factor CTCF. Considering the performance of Epitome in predicting the binding positions of CTCF, we observe that the value of adding more reference data sets starts to diminish at approximately ten data sets, and that beyond this point the performance tends to saturate. Since a similar level of saturation is not reached in the other TFs, these results may serve to provide intuition for the number of cell types which may be required to achieve high accuracy in future applications of Epitome.

**Figure 3:**
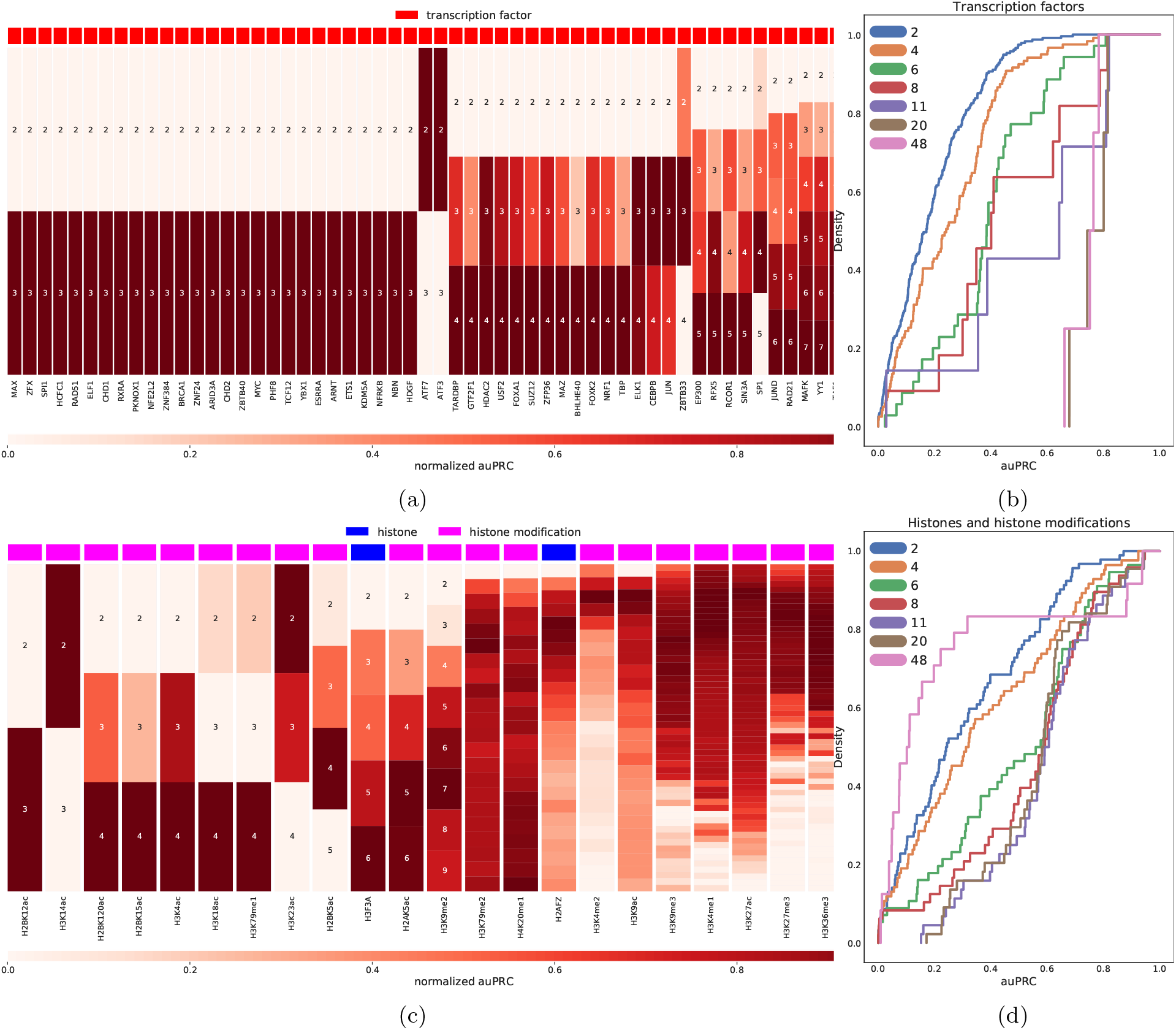
The number of cell types selected to train an Epitome model changes predictive performance of transcription factors, histones, and histone modifications. (a) Normalized mean auPRC of 59 transcription factors in heldout chromosome 7 as more cell types are incorporated into Epitome for training. For each set of reference cell types considered for a given TF, mean auPRC was calculated across four models with different combinations of training and validation cell types. y-axis indicates the addition of more reference cell types used to train each target specific model, where numbers indicate the number of cell types used to train a model. The number of training cell types considered ranges from 2 to 48 cell types. This range is dependent on the availability of reference cell types for a given transcription factor in ENCODE. (b) Cumulative distribution function (CDF) of auPRC performance of 59 transcription factors in heldout chromosome 7. Models were trained on 2 to 40 cell types. Transcription factors evaluated are listed in (a). (c) Normalized mean auPRC of 23 histone modifications and histones in heldout chromosome 7 as more cell types are incorporated into Epitome for training. Mean auPRC was calculated across four models with different combinations of training and validation cell types. y-axis indicates the addition of more cell types used to train each target specific model, where numbers indicate the number of cell types used to train a model. The number of training cell types considered ranges from 2 to 84 cell types. This range is dependent on the availability of experiments for a given histone modification or histone in ENCODE. (d) Cumulative distribution function (CDF) of auPRC of 23 histone modifications and histones in heldout chromosome 7. Models that were trained on 2 to 48 cell types were included. Histone modification and histones evaluated are listed in (c).

Interestingly, when Epitome is applied to predict the positions of histones or histone modifications, we observe a similar trend of improvement in performance, but only up to a certain level. Similar to the TFBS prediction task, the performance of Epitome starts to saturate when more than ten data sets are used as a reference. However, we also observe a marked decrease in performance when the number of cell types that are used as reference goes beyond twenty. Taken together, these results suggest an optimal regime for the number of data sets to be used by Epitome as a reference. We note that this actual number can be evaluated by cross-validation, a utility which will become crucial as the number and diversity of ChIP-seq data sets increases.

### 2.4 Considering wide genomic contexts and multiple epigenetic signals to compute cell type similarity improves the performance of Epitome

Epitome uses the chromatin accessibility similarity vector (CASV) to estimate the extent to which a query cellular context is similar to the reference cell types at each candidate locus. The CASV accounts for similarities at several scales, from a small window of 200bp around the locus in question, up to a window of size 12kbp. We next explored whether accounting for multiple levels of resolution aids in learning better decision rules. To this end, we evaluated the performance of Epitome in predicting each of the 82 epigenetic events, shown in Figure 3, while varying the size of the genomic context used for the CASV. We considered five possible sizes of genomic context to be considered by the CASV, including 0bp (the identity CASV), 200bp, 1,200bp, 4,000bp, and 12kbp. For each of the models trained in Figure 3, we trained 4 additional models, each using a different genomic context ranging from 0-12kbp. These models were similarly trained using different numbers of reference data sets ranging from 2 to 30 in size, based on the availability of data sets for a given ChIP-seq target.

Figure 4(a) and 4(b) depict the overall change in pAUC and auPRC, respectively, as larger genomic contexts are considered. The most basic models, which simply tally up the number of observed peaks (i.e., do not account for accessibility data; ”identity CASV”) as a decision rule perform the poorest, thus supporting the merit of using chromatin accessibility data. Consistently, we observe slight increases in performance as the genomic context considered by the CASV increases. This observation is most easily observed in change in AUC as wider genomic contexts are considered, shown in Supplementary Figure S12(a). We find that while performance increases with the presence of a wide genomic context for models that are trained with less than 10 cell types, histone modifications in particular suffer from considering wide genomic context when models are trained on more than 10 cell types. Figures S12(c) and S12(b) show that when histone modifications are trained on more than 10 cell types, performance decreases as wider genomic contexts are considered. These observations agree with trends seen in Figure 3(c), showing that performance of histone modifications does not saturate, but degrades, with increasing information in models trained with more than 10 cell types.

**Figure 4:**
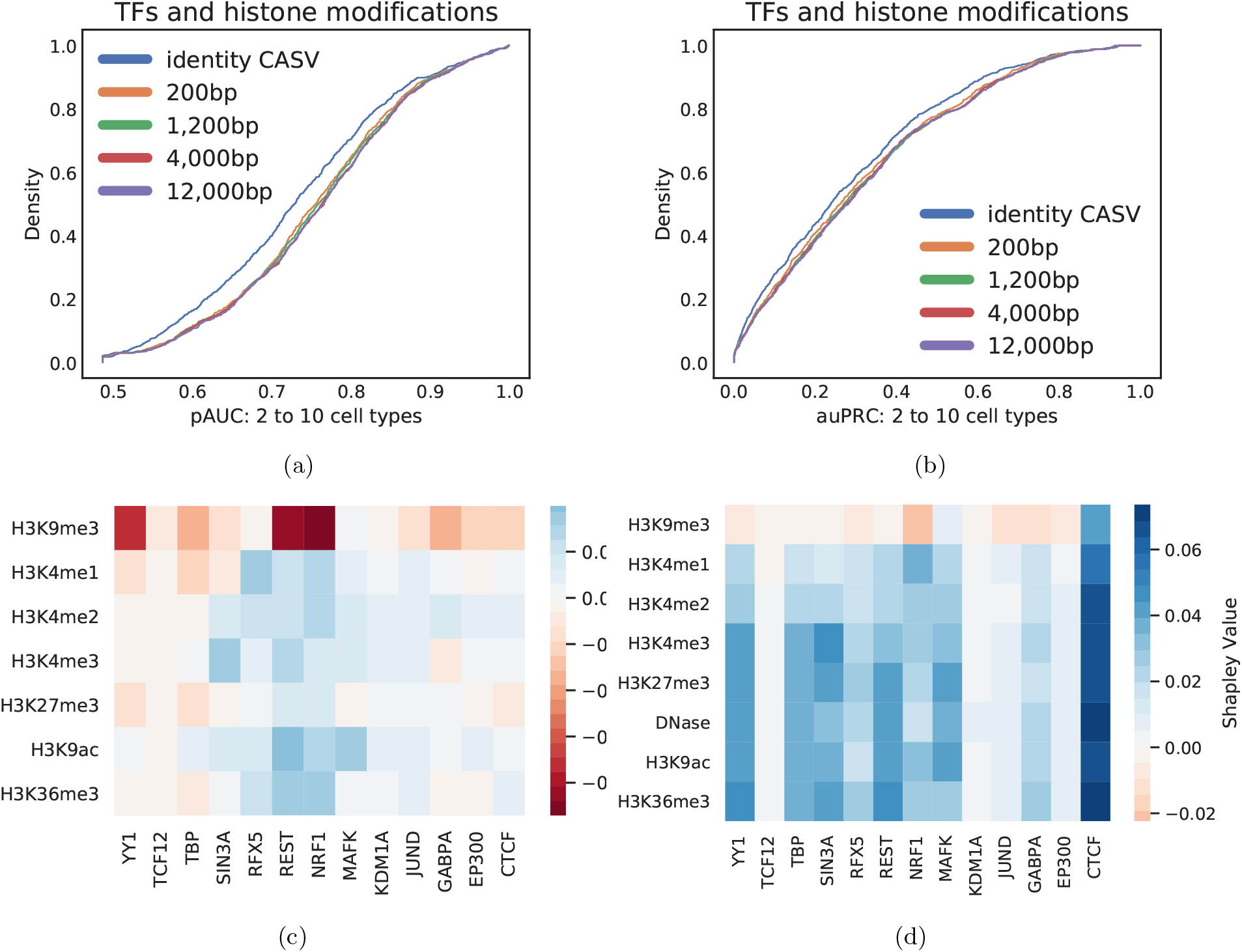
Considering wide genomic contexts and multiple epigenetic signals to compute cell type similarity improves model performance. CDFs of Epitome performance in terms of pAUC (a) and auPRC (b) as various DNase-seq window sizes are considered for computing the chromatin accessibility vector (CASV). Only DNase-seq is used to compute cell type similarity in the CASV. DNase-seq window sizes considered include the identity CASV, 200bp, 1,200bp, 4,000bp, and 12,000bp around a peak of interest. Only models training on less than 10 cell types were considered. Identity CASV implies that no CASV is used in Epitome. (c) Difference in auPRC for 13 TFs when Epitome uses a single histone modification and DNase-seq in the CASV, compared to performance when only DNase-seq is used in the CASV. All 200bp regions on chromosome 7 that have at least one ENCODE epigenetic event were evaluated, where positive include 200bp regions that overlap a ChIP-seq peak for the ChIP-seq target evaluated, and negative regions are all 200bp regions not overlapping a ChIP-seq peak. (d) Shapley values of seven histone modifications and DNase-seq demonstrating their contribution of auPRC performance of 13 TFs when incorporated into the CASV.

In all results shown thus far, Epitome only incorporates chromatin accessibility in the CASV to compute similarity between a query cellular context and reference cell types used for training. However, additional assays, such as ChIP-seq for histone modifications, are often available in addition to chromatin accessibility, and could be used to compute a more robust measure of similarity of a local chromatin environment. Histone modifications in particular provide valuable information to compute chromatin similarity, as it has been observed that DNase-seq hypersensitivity sites that are common to many cell types are only weakly correlated with certain histone marks [39]. In these cases, weakly correlated histone marks can provide somewhat independent and potentially more specific information of similarity in regions that already have similar chromatin accessibility. This is also a common use case that a query cellular context has been partially characterized with multiple experiments, including chromatin accessibility and various histone modifications (Supplementary Figure S3).

To make use of all assays that may partially characterize a cellular context, we explored whether extending the CASV to include histone modifications in addition to chromatin accessibility could improve predictive accuracy of Epitome. To test this, we evaluated the effect of incorporating various histone modifications in the CASV, and how this alteration changed the auPRC performance for thirteen TFs, listed in Figure 4(c). These thirteen TFs were selected in particular because they were available in the four reference cell lines that had measurements for seven histone modifications, allowing us to extend the CASV to use histone modifications from reference cell types. The seven histone modifications incorporated into the CASV included H3K9ac, H3K4me3, H3K4me2, H3K4me1, H3K36me3, H3K27me3, and H3K27ac.

Incorporating these seven histone modifications, as well as DNase-seq, in the CASV resulted in 256 configurations of Epitome that used all possible combinations of the seven histone modification and DNase-seq to compute the CASV. For each configuration, we trained four separate models using three reference cell lines each, and then evaluated each model on a fourth held out cell line. Final auPRC performance for each configuration was calculated across predictions for all four evaluated models. Details on how the CASV is extended to include histone modifications is described in Methods (4.3.6).

Figure 4(c) shows the difference between auPRC performance of Epitome using a single histone modification as well as DNase-seq in the CASV and auPRC performance using only DNase-seq in the CASV. A majority of these thirteen TFs see minor improvement in auPRC performance when including a histone modification, with the exception of YY1, TCF12, and TBP. All histone modifications evaluated have some positive improvement in auPRC, with the exception of H3K9me3. This observation is consistent with the association of H3K9me3 with the formation of transcriptionally silent heterochromatin.

We next sought to understand the relative contributions of DNase-seq and the seven histone modifications in the CASV towards model performance for the 13 TFs evaluated in Figure 4(c). We computed the Shapley value for each combination of experiments used in the CASV for each of the 13 TFs. The Shapley value indicates the marginal contribution computed from all possible subsets [40]. These values ultimately can highlight which experiments provide the maximal information for predicting TFBS, and which experiments should be prioritized when partially characterizing a cellular context of interest. Shapley values are shown in Figure 4(d) for each histone modification and DNase-seq, and their effect on each of the 13 TFs evaluated. On average, repressive mark H3K9me3 gives the least information when incorporated into the CASV, and in most cases, provides negative contribution to auPRC performance. We hypothesized that H3K9me3 gave the least amount of information because its correlation with TFs was low, providing the model with little to no consensus information. To test this hypothesis, we computed the Jaccard index between TFBS in each 200bp window for each TF in each training cell line and each peak indicating a histone modification used in the CASV. Indeed, H3K9me3 had the smallest mean Jaccard score across training cell lines (0.0008), compared to other histone modifications and DNase-seq (0.04 mean). These results suggest that including histone modifications in the CASV can improve predictive performance when it correlates with the epigenetic signal of interest.

### 2.5 Epitome recapitulates changes in H3K27ac over neural induction of human pluripotent stem cells

We next evaluated Epitome’s ability to leverage changes in ATAC-seq to detect changes in the acetylation of H3K27 (H3K27ac) over neural induction of human pluripotent stem cells (hPSCs). This analysis can demonstrate how well Epitome is able to leverage changes in chromatin accessibility from the same starting population to detect gradually accumulating H3K27ac marks. Temporal analysis of neural induction from hPSCs has shown that changes in chromatin accessibility precede H3K27ac, a histone mark indicative of transcriptionally active regions [41]. We therefore sought to use Epitome to predict H3K27ac in seven timepoints of neural induction by using chromatin accessibility to compare timepoints to reference cell types used to train a model.

In previous analyses of neural induction, Inoue et al. collected H3K27ac and ATAC-seq across seven time points of neural induction starting from hPSCs. Genomic regions that were enriched for H3K27ac over timepoints were grouped into six clusters, each associated with a different temporal pattern. These clusters identified H3K27ac peaks present in early induction (clusters 1-3), mid induction (cluster 4), and late induction (clusters 5-6). To determine whether Epitome could leverage ATAC-seq to identify changes in H3K27ac over neural induction, we used ATAC-seq from each time point as input into the CASV to predict H3K27ac peaks. Although the Epitome model primarily uses DNase-seq to compute the CASV and train its models, we show in Supplementary Figure S14 that Epitome can accurately predict histone modifications and TFBS when using ATAC-seq during evaluation in the CASV when Epitome has been trained using DNase-seq. For this reason, we hypothesized that Epitome could provide sensitive predictions of H3K27ac by using the CASV to compare similarity between DNase-seq in reference cell types and ATAC-seq from neural differentiation time points. We trained an Epitome model to predict H3K27ac peaks using all ENCODE reference cell types that had both H3K27ac and DNase-seq, resulting in 15 reference cell types.

Here, we compare Epitome to an additional benchmark method specifically designed to predict histone modifications, called DeepHistone [42]. DeepHistone is a deep learning method for predicting seven histone modifications, including H3K27ac, and uses DNA sequence and chromatin accessibility as features to provide predictions specific to a cellular context. For further insight we also added a simple baseline predictor, which uses enrichment signal from ATAC-seq peaks to directly indicate the signal of H3K27ac (i.e., predict an H3K27ac peak whenever there is an ATAC-seq peak). We chose this baseline because increased acetylation of H3K27 is often observed in accessible regions, and in neural induction it was often observed to be preceded by opening of the chromatin [41]. Figure 5(a) shows the ROC and PR curves for H3K27ac peaks across seven time points in 39,000 regions for the three comparative methods. All autosomal regions that had an H3K27ac peak in at least one of the seven timepoints were considered. For all time points, Epitome performs significantly better than the baseline predictor as well as DeepHistone, with a gain of mean auPRC of 0.09 and 0.21, over the baseline ATAC-seq predictor and DeepHistone, respectively. This boost in performance seen by Epitome suggests that using known H3K27ac peaks from ENCODE cell types, along with the CASV to inform the model of similarity between ENCODE cell types and differentiated hPSCs, improves predictive performance of H3K27ac predictions over baseline predictors that do not use signal from the epigenetic event of interest as features.

**Figure 5:**
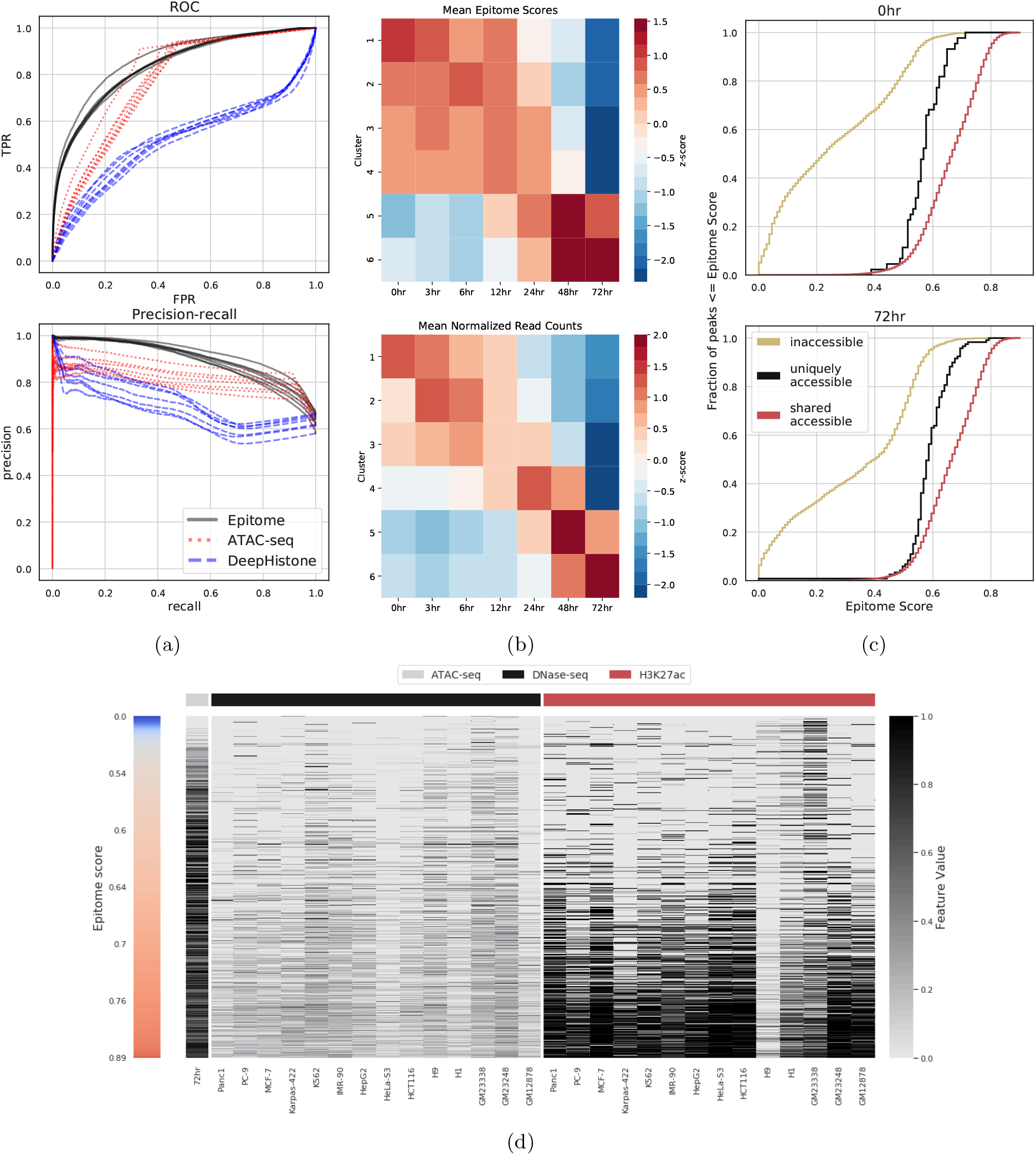
Epitome detects differential H3K27ac across seven time points in neural differentiation from human pluripotent stem cells. (a) ROC and PR curves for predictions of H3K27ac peaks using three methods: Epitome, an ATAC-seq enrichment baseline predictor, and DeepHistone, at seven time points of neural differentiation. (b) (Top) Mean Epitome scores of H3K27ac peaks across seven time points in six clusters from 2,400 temporal peaks [41]. (Bottom) Mean normalized H3K27ac read counts across seven time points in six clusters. Rows are standardized. (c) CDFs of Epitome scores for H3K27ac peaks at 0hr and 72hr in regions that are uniquely accessible to a timepoint (black), are inaccessible for a timepoint (yellow) and have shared accessibility across all timepoints (red). Both 0hr and 72hr timepoints have low scores in inaccessible regions. Regions of unique and shared accessibility have and moderate and strong scores, respectively. (d) Heatmap of features used by the Epitome model for 25,762 genomic regions containing H3K27ac peaks at 72hr. Black and red columns indicate values of DNase-seq and H3K27ac features from reference cell types. ATAC-seq column, labeled in grey, indicates presence of absence of ATAC-seq peaks in the 72hr time point. Color bar on left represents Epitome scores, where blue represents instances of false negatives and red represents instances of true positives.

Although Epitome can predict H3K27ac peaks better than baseline predictors and existing models, ROC and PR curves could not illuminate whether Epitome could identify H3K27ac peaks that were unique to each time points. We therefore sought to show that Epitome can leverage changes in chromatin accessibility over time to detect time point specific H3K27ac peaks. Figure 5(b) demonstrates the mean predictions of Epitome (top) and the normalized H3K27ac read counts (bottom) in six H3K27ac clusters indicating key H3K27ac peaks of early, mid, and late induction across 2,400 genomic regions. These six clusters were defined by Inoue et al [41]. Epitome predictions show that at 48hr and 72hr, there is an increase in H3K27ac in clusters 5 and 6. Additionally, clusters 2, 3, and 4 have increased H3K27ac between time points 3hr to 24hr, and decrease at 72hr. These broad trends of increased H3K27ac in late induction clusters 5 and 6 and in early to mid induction clusters 2-4 agree with H3K27ac normalized read counts shown in the bottom heatmap.

In this analysis, Epitome leverages ATAC-seq to predict time point specific H3K27ac. This means that for a given region of the genome, Epitome will predict the same probabilities for H3K27ac for all samples, unless ATAC-seq signal varies across samples in or around a given region of interest. Because of this, we would expect Epitome to provide sensitive results when predicting H3K27ac in regions with differential ATAC-seq between time points. To assess this hypothesis, we evaluated Epitome H3K27ac predictions at 0hr and 72hr, and separated predicted regions into three groups: regions accessible in all time points, regions only accessible in either 0hr or 72hr, and regions not accessible in either 0hr or 72hr. Figure 5(c) shows the CDFs of Epitome H3K27ac predictions at 0hr and 72hr across these three groups. At both the 0hr and 72hr time points, Epitome predictions are higher in accessible regions, compared to inaccessible regions. These results suggest that Epitome is effectively leveraging accessibility to predict H3K27ac. We note that for both the 0hr and 72hr predictions, Epitome is able to easily detect H3K27ac in shared accessible regions of the genome, compared to regions of the genome that have differential accessibility in either the 0hr or 72hr time points. In regions of differential accessibility, Epitome predictions are lower than those found in regions of shared chromatin accessibility. This is most likely an artifact of decreased availability of evidence for H3K27ac in the training cell types in regions that have differential accessibility. In regions of shared accessibility, training cell types have H3K27ac marks in a median of 8 out of 13 cell types, compared to peak regions unique to either 0hr or 72hr time points, which have H3K27ac marks in a median of 2 out of 13 cell types. Thus, commonly accessible regions have greater evidence of H3K27ac marks in training cell types, and are predicted with greater confidence.

Although Epitome can predict H3K27ac marks better than existing methods, Epitome still incorrectly identifies a subset of peaks as false negatives, and a subset of non-peak regions as false positives. Figure 5(d) and Supplementary Figure S13(a) show Epitome predictions for H3K27ac at 72hr in peak and non-peak regions respectively. Figure 5(d) demonstrates that while a majority of H3K27ac marks are correctly identified, there is a small subset of peaks that are incorrectly identified as false negatives. Additionally, many of the non-peak regions are identified as peaks. Because of this trend, we sought to identify the source of the false positives and negatives within the context of the reference cell types used for training. Figure 5(d) shows reference data sets used as input in the Epitome model for predicting H3K27ac marks in peak regions at 72hr. Epitome’s true positives are shown towards the bottom of the heatmap, while false negatives are displayed at the top. This plot shows that true positives have ATAC-seq accessibility at the 72hr time point, as well as supporting H3K27ac peaks from the reference cell types used for training. False negatives generally have less ATAC-seq accessibility at 72hr, and less H3K27ac peaks from the reference cell types. These results demonstrate that false negatives were generated from regions that had little to no accessibility at the 72hr timepoint, as well as little support for H3K27ac in reference cell types. This trend is similarly shown in the predictions of non-peak regions at 72hr, shown in Supplementary S13(b), where false positives, shown towards the bottom of the heatmap, have more accessibility at 72hr and representation of H3K27ac peaks in the reference cell types than peaks correctly identified as true negatives. These results show that false positives and false negatives result from unexpected patterns in chromatin accessibility and H3K27ac in both the query cellular context and reference data sets.

## 3 Discussion

We presented Epitome, an algorithm to predict the probability of observing enriched genomic regions for transcription factor occupancy and histone modifications in a new cellular context, by estimating similarities to other cell types in which epigenetic measurements are already available. Due to Epitome’s design, both the quality of predictions and quantity of epigenetic events that can be predicted will be further improved and expanded as more ChIP-seq targets are measured in more cell types.

Existing data consortia can expand their repertoire of regulatory regions through accumulation of ChIP-seq targets in two ways, both of which uniquely affect the performance of Epitome. First, regulatory regions for a specific ChIP-seq target can be further annotated through the accumulation of additional ChIP-seq targets (primarily transcription factors) in already well characterized cell lines and primary cell subsets. This accumulation strategy has been exercised in ENCODE 3 [1], and is demonstrated through the accumulation of DNA-associated proteins in the already well annotated K562 and HepG2 cell lines. This accumulation of ChIP-seq experiments in well annotated cell lines provides Epitome with additional data to more accurately predict ChIP-seq targets as their coverage across cell types increases. The benefits of this type of data expansion in Epitome is demonstrated in Figure 3, which shows that the predictive performance of many ChIP-seq targets increases as their characterization across reference cell types increases. Secondly, data consortia can expand their annotation of regulatory elements that are highly cell type selective. This requires the expanded annotation of regions in cell types and conditions that have rare, condition specific regulatory elements. This particular type of expansion is going to be a key effort in ENCODE 4 [43] through the expansion of the collection of cellular contexts in which the experiments are conducted.

Because Epitome leverages cell type similarity to predict epigenetic events in a new cellular context, Epitome can only make predictions in regions that were previously annotated by reference datasets for an event of interest. This limitation is demonstrated in Figure 5(d), where Epitome fails to correctly identify H3K27ac in regions that are not already annotated in the reference cell types used for training. In these cases, Epitome forfeits sensitivity in regions that are not annotated in reference cell types. However, previous work has demonstrated that as more cell types are characterized, the number of unique events discovered in a new cell type diminishes [29, 27]. Figure 1(a) similarly shows that across 152 ChIP-seq and DNase-seq experiments, the fraction of unique peaks discovered in each cell type used in Epitome decreases as more cell types are characterized for a given experiment. These trends show that although Epitome cannot predict epigenetic events in all regions of the genome, these regions have diminishing chances of containing unique regulatory elements as more cell types are collected by large data consortiums and are incorporated into Epitome. Additionally, further initiatives such as ENCODE 4 [43] will increase the coverage of regulatory regions that are cell type selective.

When considering various design decisions in Epitome, in particular the construction of the CASV, we looked to ChIP-seq centric studies to assess the best way to calculate cell type similarity. Because of the wealth of DNase-seq data available in the public domain across numerous cell lines and primary cells [1, 27], we have focused on using DNase-seq as the primary method to compute similarity between cell types. However, we additionally evaluated the frequency at which ChIP-seq targets were present in studies in ChIP-Atlas that contained at least two ChIP-seq experiments. These results are shown in Figure S3, and show that many histone modifications, including H3K27ac, H3K27me3, and H3K4me1 and H3K4me2, are frequently collected, along with at least one other additional ChIP-seq target of interest. Excitingly, Figure 4(d) shows that H3K4me1 and H3K4me2 provide the greatest information gain, in terms of their expected added value (Shapley value), when incorporated into the CASV to predict TFBS. These results suggest that Epitome is well situated to utilize this wealth of histone modification data in personal studies to gain more confident predictions of TFBS.

Lastly, we acknowledge the distinction between models that are designed to predict a single epigenetic event and multiple epigenetic events in the same model in held out cellular contexts. As shown in Figure 2 and Supplementary Figure S6(b), models that are trained individually on epigenetic events of interest often outperform models trained jointly. While existing literature in the machine learning community observes this phenomenon [33], it has not been acknowledged within the realm of methods for predicting epigenetic events. Going forward, this loss in performance that is accompanied with choice in model design should be considered, or reconciled with the utilization of methods that determine which epigenetic events can be trained jointly without loss of performance [33].

Epitome is an open-source project, and is available on GitHub and can be installed using the Python Package Index (PyPI). We believe that Epitome will have a substantial impact on studies that have collected data that provide a limited view of the epigenome through chromatin accessibility. In particular, Epitome can provide a mechanistic complement to methods that annotate accessible regions based on enrichment of nearby genes [37]. Such methods provide insight into which genes are affected by changes in the epigenome at cis-regulatory regions. However, Epitome can analyze TF activity and histone modifications of trans-regulatory regions independently of distance to nearby genes. Utilizing existing cis-regulatory gene-ontology based approaches, along with Epitome, can provide comprehensive annotation of cis- and trans-regulatory regions for any given dataset partially characterized with chromatin accessibility.

## 4 Materials and Methods

### 4.1 Processing ENCODE data for model training and validation

Epitome uses peak called DNase-seq and ChIP-seq from ENCODE for training, validation, and testing models. Peak called DNase-seq and ChIP-seq from the hg19 and hg38 genome were processed from the ENCODE portal (version v94) [2]. We utilized ChIP-seq and DNase-seq experiments from all cell types that had available DNase-seq and contained ChIP-seq peaks for at least 1 ChIP-seq target. Only ChIP-seq targets that had experiments for more than three cell types were considered. We only considered experiments that were compliant with the ENCODE pipeline and had no audit errors. This filter resulted in 17 cell types with 153 unique ChIP-seq targets for hg19, and 93 cell types with 250 unique ChIP-seq targets for hg38. We used optimal irreproducible discovery rate (IDR) peaks for ChIP-seq experiments when available in the ENCODE portal and MACS2 called peaks for the remaining DNase-seq experiments [44, 45]. All ENCODE accessions used are listed in Supplementary Table S2.

We next joined all peak sets together in 200bp binned regions using bedtools [46]. This join resulted in 8.5 million binned regions for h19, and 10.9 million regions for hg38, each of which contained at least one DNase-seq or ChIP-seq peak in at least one of the included cell types. Mitochondrial and sex chromosomes were removed from the peak set. The final regions were divided into a set for training, validation, and test. The validation set contains all 200bp regions on chromosome 7 that have at least one DNase-seq or ChIP-seq peak. The test set contains all 200bp regions on chromosomes 8 and 9 that have at least one DNase-seq or ChIP-seq peak. The training set consists of all remaining autosomal chromosomes, including chromosomes 1-6, and 10-22.

Code for downloading and processing ENCODE [2] DNase-seq and ChIP-seq datasets used for Epitome training and validation can be found in the Epitome GitHub repository. Curated datasets for hg19 and hg38 are publicly available in Amazon S3.

### 4.2 Processing ChIP-Atlas database for analyzing the fraction of unique peaks across cell types for available ChIP-seq targets

In Figure 1(a), we calculate the fraction of unique peaks found in a held out cell type for a given ChIP-seq target, or DNase-seq. We use ChIP-seq and DNase-seq from ChIP-Atlas [35], a comprehensive database of ChIP-seq and DNase-seq experiments that are publicly submitted in Sequence Read Archives (SRA) [47]. We selected all ChIP-seq targets from ChIP-Atlas that were observed in at least three cell types, and selected all cell types that had DNase-seq and at least one ChIP-seq experiment. This filter resulted in downloading experiments across 197 cell types for 694 ChIP-seq targets for hg19 and 194 cell types for 697 ChIP-seq targets for hg38. Experiments used from ChIP-Atlas are listed in Supplementary Table S3. Significant peaks from each experiment were defined as peaks called from MACS2 [45] with a Q-value < 1.0e − 5. Because many of the cell type and ChIP-seq target combinations in ChIP-Atlas have multiple replicates, we used a consensus approach to merge peaks from replicates. This consensus approach chose all peaks for a given type/target combination that were present in at least 25% of the replicates. If only 2 replicates were available, all peaks were used. If only 3 replicates were available, all peaks present in at least two replicates were used. Final consensus peaks for each type and target combination were binned into 200bp regions of the genome.

In Figure 1(a), we calculate the fraction of unique peaks that are observed in a held out cell type in ChIP-Atlas, compared to peaks identified in remaining cell types for 121 ChIP-seq targets that had experimental data from at least four cell types. Let *A* be the set of peaks in a held out cell type for a ChIP-seq target of interest, where this ChIP-seq target is available in *n* cell types. Let *B_i_* be the set of all peaks combined from a set of randomly selected cell types *i*, where |*i*| < *n*.

We then calculate the fraction of unique peaks shown in Figure 1(a) and Supplementary Figure S2 as:

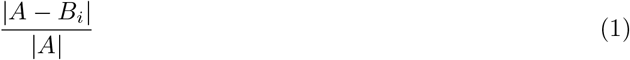

Ideally, for each ChIP-seq target, we would calculate the fraction of unique peaks for every held out cell type, for every combination of cell types in which that ChIP-seq target is measured in. However, for some ChIP-seq targets, this is computationally infeasible. For a target available in *n* cell types, there are 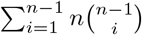 combinations of held out cell types and choices of reference cell types. For example, H3K4me3 is available in 138 cell types, resulting in 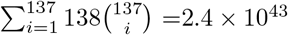 combinations. Due to this constraint, we randomly choose a held out cell type to define *A*, and sample the remaining cell types to define set *B_i_* for a given target. |*i*| is chosen randomly for each sample, and ranges from 1 to *n* − 1 for a given target and held out cell type combination. We perform this sampling procedure 3,000 times to calculate the fraction of unique peaks for 3,000 combinations of cell types chosen to form *B_i_* for a given ChIP-seq target.

Because we want to observe general trends in the relationship between the fraction of unique peaks and number of available cell types, we do not want certain ChIP-seq targets available in many cell types to overweight the results shown in Figure 1(a). Therefore, for each number of cell types, shown on the x-axis in Figure 1(a), we calculate the weighted mean and weighted standard deviation of fraction of unique peaks for all combinations of ChIP-seq targets and held out cell types. Each data point representing a ChIP-seq target and held out cell type is weighted inversely proportional to the number of samples for that TF observed at the number of selected cell types.

### 4.3 Overview of Epitome

Epitome predicts the genomic locations of epigenetic events that can be measured through ChIP-seq, including protein-DNA binding sites, histone modifications, chromatin modifier binding sites, and locations of histone variants. All ChIP-seq targets and cell types used in Epitome are listed in Supplementary Table S2. Epitome can predict unmeasured epigenetic events in any cellular context of interest, as long as genome-wide measurements of chromatin accessibility in that context is available. This is shown in Figure 1(b), where chromatin accessibility from a query cell type *c′* is compared to chromatin accessibility from reference cell types in order to predict epigenetic signal in *c′*. As input, Epitome requires peak called DNase-seq or ATAC-seq in the form of bed or narrow peak file formats from the query cellular context of interest, as well as a similar formatted file containing all genomic regions to be queried. For training, Epitome additionally requires a set of cell types that have both measured epigenetic signal for the epigenetic event of interest and chromatin accessibility. We refer to these cell types with known epigenetic signal and available chromatin accessibility as reference cell types. Because of the abundance of DNase-seq available in ENCODE [34], Epitome uses DNase-seq as the primary measure of chromatin accessibility for model training. However, Epitome can make predictions in a query cellular context using other assays that measure chromatin accessibility, such as ATAC-seq (see Supplementary Figure S14). We leverage chromatin accessibility from all reference cell types to compute an explicit metric of similarity between reference cell types and the query cellular context. This metric is referred to as the chromatin accessibility similarity vector (CASV), and compares the similarity between reference cell types and a query by comparing the similarity of chromatin accessibility at multiple windows of resolution surrounding a genomic locus of interest (See Methods 4.3.1). The CASV, along with binary epigenetic events from reference cell types, are used as features in a neural network, which uses the CASV to weigh the importance of each reference cell type for predicting epigenetic events in the query. Final predictions are the probabilities of observing each epigenetic event in question.

The set of candidate positions on which Epitome is applied consists of all loci in which the epigenetic event of interest has been observed in at least one of the reference cell types. This set of candidates is determined by unifying all the observed peaks from reference cell types (Methods 4.1). One important observation is that Epitome simplifies the input epigenetic data as binary, and the features it uses represent the presence or absence of ”peaks” (or a signal). While this can lead to loss of information, it helps mitigate bias and noise arising from variations in sequencing depth, quality of antibodies and other technical factors that may affect the data quantitatively [48]. This strategy also facilitates the use of multiple types of epigenetic events as features (in addition to the mandatory accessibility data) without the need for calibration of their dynamic range or other quantitative attributes. Specifically, if a query cellular context has available measurements of certain histone modifications in addition to chromatin accessibility, these can be used to better evaluate similarity to the reference cell types and thus guide the prediction process (by extending the definition of CASV; see Methods 4.3.6).

Underlying Epitome is a feed forward neural network (NN). Epitome’s underlying NN can be written as *P*(*y*|*x*, **w**). For the problem of predicting the presence of ChIP-seq peaks, *y* ∈ [0, 1] is a set of classes, where each class is a ChIP-seq target predicted by Epitome. The values of *y* indicate the presence (1) or absence (0) of a ChIP-seq peak in a given 200bp region in the genome. **w** are a set of parameters learned by the Epitome model. *x* represents a set of features used to train an Epitome model. Features *x* contain ChIP-seq peaks from well characterized ENCODE cell types, as well as a measure of chromatin accessibility similarity between ENCODE cell types and query cellular context *q*. These features *x* are explained in greater detail in Sections 4.3.2 and 4.3.1. Figure 1(b) represents a schematic of Epitome. This figure visualizes how Epitome computes the similarity of chromatin accessibility between ENCODE cell types and *q* and uses this similarity as input into the model, along with binarized ChIP-seq peaks from ENCODE cell types, to predict peak probabilities for ChIP-seq targets in *q*. Model output are the probabilities of observing a peak for each ChIP-seq target being predicted in *q* at a 200bp genomic region *i*. Epitome can predict individual or multiple ChIP-seq targets in a single model, depending on which ChIP-seq targets and cell types are selected to train the model. We model the loss as the sigmoid cross entropy between the model’s prediction of ChIP-seq targets 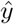 and the ground truth labels *y*. Cross entropy loss is defined in defined in Equation 2.

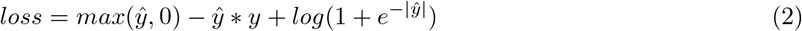

We parameterize the model, *P*(*y*|*x*, **w**), by constructing cell type specific channels for each reference cell type used in training. Cell type specific channels are visualized in Figure 1(b) and explained in Section 4.3.2.

#### 4.3.1 Measuring cell type similarity with the Chromatin Accessibility Similarity Vector (CASV)

As previously mentioned, Epitome uses a metric of cell type similarity to weigh the importance of reference cell types when predicting epigenetic signal in a query cellular context. This metric is referred to as the chromatin accessibility similarity vector (CASV). Figure 1(b) visualizes how the CASV is calculated between a query cellular context *q* and a reference cell type *k* at a 200bp genomic region *i*. The CASV compares the agreement of binarized accessibility peaks between *q* and *k* at varying genomic windows up to 12kbp surrounding *i*. This maximum genomic distance of 12kbp was determined through hyperparameter search using validation chromosome 7 (See Figure S12(a)). Epitome uses the CASV to determine how similar *q* is to all reference cell types. This similarity is leveraged to determine the relative importance of each reference cell types in predicting ChIP-seq peaks for *q*. Without the CASV, Epitome is not able to provide cell type specific predictions for *q*, as shown in Supplementary Figure S12(a).

We formally define the CASV in Equation 3. *a^k^* and *a^q^* indicate binarized chromatin accessibility peaks for cell types *k* and *q* binned in 200bp bins across the genome, where 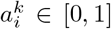 represents the presence (1) or absence (0) of a peak in bin *i*. The CASV calculates the fraction that *a^k^* and *a^q^* have shared chromatin accessibility peaks in region *i*. It also calculates the fraction that *a^k^* and *a^q^* have shared chromatin accessibility peaks in larger genomic windows surrounding region *i*. We consider exclusive 200bp genomic windows surrounding *i* in windows *R* = {*r_z_*; 0 ≤ *z* ≤ 3}. We set |*r*_0_| = 1, |*r*_1_| = 5, |*r*_2_| = 19, and |*r*_3_| = 59, representing the number of 200bp bins considered in each window surrounding *i*. We first compute *CASV_n_*,a vector of fractions of how many bins *a^k^* and *a^q^* agree for each window *r_z_* ∈ *R* surrounding region *i*, relative to the size of each window. Here, agreement criteria is met if both *q* and *k* have a peak or do not have a peak. Each window *r* is computed exclusively from smaller windows to avoid redundancy:

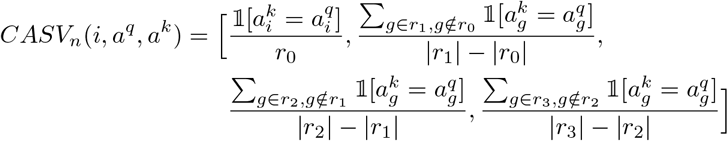

Because chromatin accessibility peaks are sparse across the genome for most cell types, regions where both *q* and *k* have a chromatin accessibility peak are rare. Therefore, high values of *CASV_n_* mostly represent regions of the genome where *a^q^* and *a^k^* do not have peaks. However, to gain a complete understanding of cell type similarity, we must also know where *a^q^* and *a^k^* are both accessible. We encapsulate shared accessibility in *CASV_p_*. This metric indicates shared regulatory regions between cell types [13].

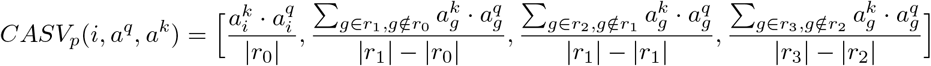

The final *CASV* for a region *i* is a concatenation of the agreement and positive *CASV* vectors:

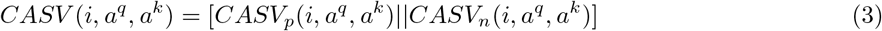

*CASV*(*i, a^q^, a^k^*) are used as features in the Epitome model, described in Equation 4. The final *CASV* comparing similarity between two cell types is a vector of length 8. Using the CASV, Epitome is thus able to learn different weights corresponding to elements in the CASV, and can thus determine the relative importance of chromatin accessibility similarity at varying distances from the genomic region of interest for each reference cell type.

In practice, Epitome uses binarized peaks called from DNase-seq to calculate the CASV during model training. However, ATAC-seq can also be used to compute the CASV (See Supplementary Figure S14). In the case of model training, the CASV measures similarity of binarized DNase-seq peaks between a held out cell type *k′* and each training cell type *k* ∈ *C, k* ≠ *k′*, where *C* is the set of reference cell types in which the epigenetic event of interest has been measured. In the case of evaluating a query cellular context *q*, the CASV measures either binarized DNase-seq or ATAC-seq peak similarity between *q* and each reference cell type *k* ∈ *C*.

#### 4.3.2 Constructing features from ENCODE cell types

Epitome trains on multiple reference cell types in a single model. These reference cell types are primarily taken from ENCODE. This allows Epitome to jointly learn from multiple cell types, without biasing models to overfit to a single training cell type. Therefore, features *x* for training a model contain cell type specific features for each cell type *k* ∈ *C* that is selected for training. Here, we notate the number of cell types used to train a model as *n*, which is equivalent to |*C*|. Epitome can train on 2 to 93 ENCODE cell types in a single model. Because each reference cell type is uniquely characterized with different ChIP-seq targets, not all cell types can be utilized by Epitome to predict binding for a ChIP-seq target of interest. Therefore, the cell types chosen to train a model is determined by the number of cell types that have available ChIP-seq experiments for the ChIP-seq targets of interest. Figure 1(b) demonstrates an example schematic of an Epitome model in which three reference cell types are used for training a model that predicts an epigenetic event for a single ChIP-seq target.

The set of cell type specific features *x^k^* for a cell type *k* selected for training is written in Equation 4. In Equation 4, 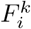 represents binarized ChIP-seq peaks in a 200bp region *i* for the set of all ChIP-seq experiments that are available from cell type *k*. *CASV*(*i, a^q^, a^k^*) is the chromatin accessibility similarity vector (CASV, see Section 4.3.1), and measures the similarity between chromatin accessibility in the query cell type, *a^q^*, and the chromatin accessibility in the training cell type, *a^k^*, at region *i*. ∥ represents concatenation.

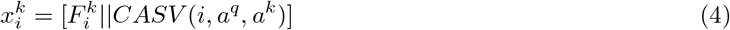

Finally, 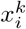 is input into a cell type specific channel with two densely-connected layers. The input dimension for each cell type channel is the number of ChIP-seq targets included in the model and available in *k*, plus the dimension of the CASV(See Equation 3). This dimension can range from 9 (1 ChIP-seq target + 8 dimensional CASV) to 258 (the maximum number of ChIP-seq targets in a dataset + 8 dimensional CASV). The dimension of the first layer is the input dimension divided by 2. The dimension of the second layer is the input dimension divided by 4. Each layer uses a hyperbolic tangent activation function. The output of the last layer from each cell type channel is combined into a final output layer that applies a sigmoid non-linearity to gather final predictions 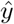.

#### 4.3.3 Training an Epitome model

Epitome trains a model on *n* ENCODE cell types by using *n*−1 cell types for features *x* (defined in Equation 4) and a held out cell type as labels *y*. Epitome can train using up to 93 cell types in a single model, depending on the data availability of cell types for the ChIP-seq target being evaluated. Training on multiple cell types allows Epitome to generalize well to new cell types, and allows Epitome to perform comparably to methods that see the cell type being evaluated during training (Supplementary Figure S10). To make full use of publicly available ChIP-seq experiments and effectively generalize to new cell types, Epitome uses a cell type rotation mechanism for training. This mechanism rotates through which cell type is used for *y*, and uses the remaining cell types to construct *x*. This rotation mechanism is repeated for all regions of the genome, excluding validation and testing regions, until the method converges. Algorithm 1 demonstrates how Epitome iterates through all training regions and all ENCODE cell types to update model parameters, using a different training cell type for *y* in each iteration. This procedure is visualized in Supplementary Figure S4, where at each iteration, the choice of cell types used in *x* and the cell type used for *y* are rotated through until the model has converged.

Note that when a given cell type is used for labels *y* during training, its cell type specific features are still included in *x* through its respective cell type specific channel. Although this allows the model to see the labels during training, it also allows the model to learn the importance of cell type similarity in predicting ChIP-seq peaks. In this case, the model will compute identity cell type similarity between the cell type used for *y* and its features in *x*. This high similarity ultimately teaches the model to up-weight features from cell types that are similar to the cell type being predicted.

We determine convergence of Epitome by defining an early stopping criterion in Methods 4.3.5 that determines when to stop training. This criteria halts training of models when validation loss no longer sees improvement. Supplementary Figure S16 visualizes changes in validation and training loss for a subset of 18 ChIP-seq targets in a joint Epitome model trained for 2000 iterations, where training loss is calculated as the mean sigmoid cross entropy across 10,000 randomly selected training regions, and the validation loss is calculated as the mean sigmoid cross entropy across all 200bp regions on chromosome 7 that have epigenetic signal from at least one of the 18 ChIP-seq targets. Epitome sets a batch size of 64, and trains between 800 and 5,000 batches.

Epitome models require different memory requirements and time allocation based on the number of ChIP-seq targets included in a model, and the number of batches required for convergence. Models trained on only 1 ChIP-seq target, with few available reference cell types take 0.5GB of RAM (16 features from reference cell types), while methods training on many ChIP-seq targets with many reference cell types, require up to 20 GB of RAM (for ChIP-seq targets resulting in 928 features from reference cell types). On hardware configurations defined in Methods 4.9, run times range from 14 minutes (for 800 batches) to 50 minutes (for 5,000 batches). Run times depend on the number of batches required for convergence.

**Algorithm 1.**
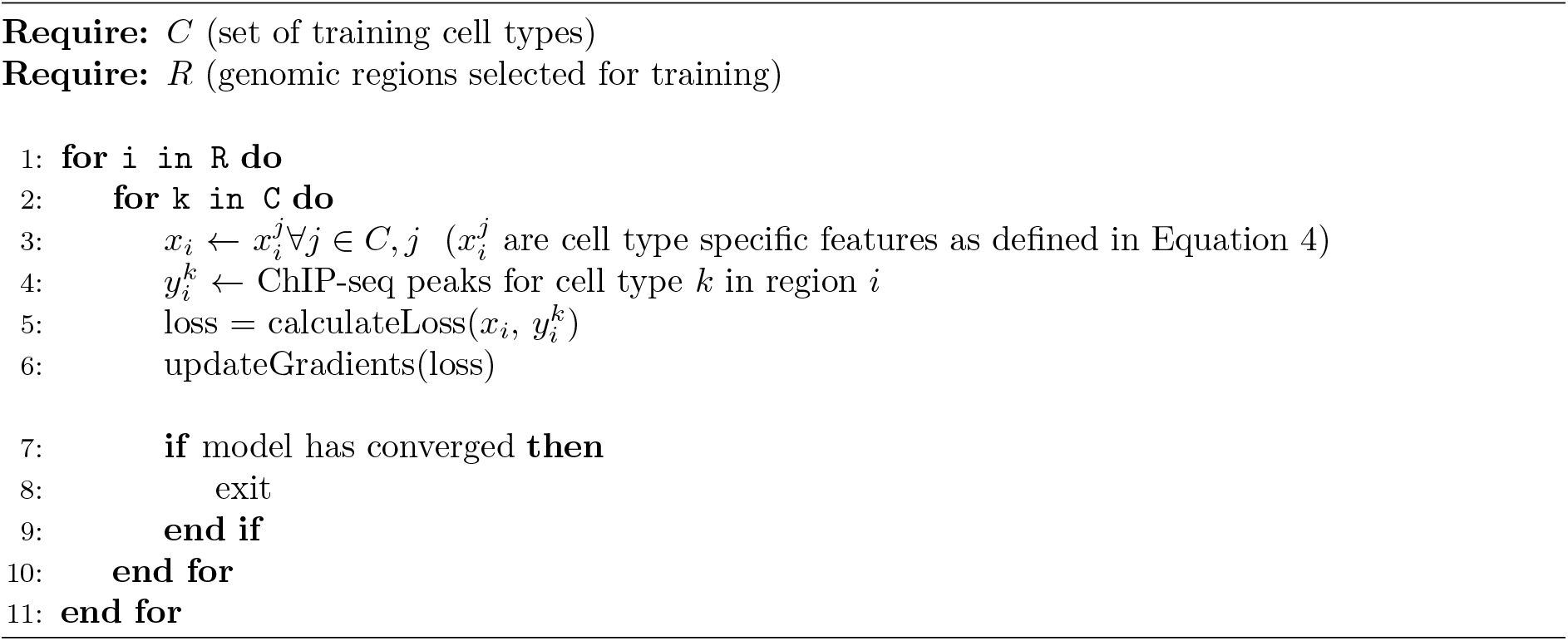
Rotate through cell types for training

#### 4.3.4 Sampling underrepresented ChIP-seq targets

When training a model for multi-label classification, imbalance across labels in the training data can result in a model that is biased towards learning over-represented labels. This problem is known as imbalanced learning. Imbalanced learning is present in models that predict peaks for multiple ChIP-seq targets because the quantity of peaks across the genome for different ChIP-seq targets is highly variable. As a solution to the problem of imbalanced learning when predicting ChIP-seq peaks, we over-sample underrepresented ChIP-seq targets to create a dataset with similar distributions of peaks for each ChIP-seq target in a multi-label model. This allows the model to see similar counts of positive instances for each ChIP-seq target during training.

To effectively over-sample underrepresented ChIP-seq targets, we borrow key incites from existing literature [49] which calculates an imbalance ratio, *IRLbl*, for each label to determine which labels need to be over-sampled. *IRLbl* measures how imbalanced a given label in a dataset is, relative to the other labels. Higher values of *IRLbl* indicate that a given label is more imbalanced, and thus needs to be over-sampled. *IRLbl* is defined in Equation 5. In this equation, *D* = (*x_i_*, *y_i_*) represents the multi-label dataset, L represents the set of labels (or ChIP-seq targets), and and *y_i_* represents labels for the *i*th instance of dataset *D*.

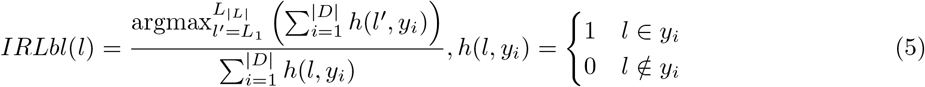

For each label *l*, *IRLbl*(*l*) can then be compared to the mean imbalance ratio, *meanIR*, for all labels *l* ∈ *L*. For each label *l*, if *IRLbl*(*l*) is greater than the *meanIR*, we re-sample *k_l_* regions in the genome that have a ChIP-seq peak for *l*. Here, we set *k_l_* to be:

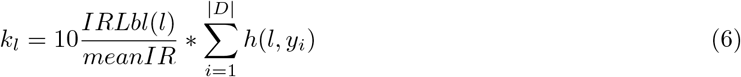

The final set of training instances includes all original instances in *D*, as well as instances oversampled for all labels *l* ∈ *L* with *IRLbl*(*l*) > *meanIR*.

In the case where Epitome is only trained on one ChIP-seq target, multi-label sampling is not required. In this case, we undersample non-peak instances for the single ChIP-seq target so that there are 10 times more non-peak instances, compared to the number of positive peak instances.

#### 4.3.5 Training with early stopping to prevent overfitting

Like all machine learning models, Epitome is susceptible to overfitting by memorizing the detail and noise of reference cell types, which can negatively impact performance on unseen cellular contexts. Although overfitting is a common problem with neural network architectures [50], regularization methods such as early-stopping methods can help [50]. Early stopping methods halt training once the model stops improving on a validation set, effectively stopping training before the model begins to overfit. Without this step, the model may memorize biological contexts specific to the cell types being trained on. Anchor, a co-winner of the 2017 ENCODE-DREAM challenge, addresses this problem by implementing a crisscross validation early stopping scheme that evaluates performance on a held out validation set during training to determine when to stop training [51]. Anchor splits its training data into two sets (train and train-validation) and computes the loss on the train-validation set after every 1000 batches of training. Once the train-validation loss plateaus, the model stops training [51].

Epitome similarly provides an optional early-stopping method that evaluates on a subset of the genomic regions to determine when to stop model training. First, genomic regions are split into two sets: a training set consisting of chromosomes 1-6,10-21 and a validation set consisting of chromosome 22 (called the train-validation set) that is used to evaluate the model during training. Epitome then trains a model on the training set with the specified training cell types and computes the weighted mean sigmoid cross entropy loss (called the train-validation loss) on the train-validation data set every train-validation iteration (200 training batches). We found that validating every 200 batches was more effective than at a lower frequency, as most of Epitome’s learning occurs in the first 1000 training batches. To speed up run-time but prevent Epitome from overfitting to the train-validation set, we sample 1000 points from the train-validation set every train-valid iteration (200 training batches) using the sampling method described in Section 4.3.4. Once the model’s train-validation loss stops decreasing, Epitome halts model training.

Because the train-validation loss might increase before decreasing to a minimum [52], we allow the model to train for up to 5 additional train-validation iterations after the train-validation loss stops improving, in order to prevent under-fitting. Sometimes the model might still over-train if the train-validation loss decreases by minuscule amounts every train-validation cycle, as the model will continue to train without significant improvements. Thus, we added a min-delta hyper parameter that requires the train-validation loss to decrease by a certain amount from the best train-validation loss in order to qualify as a train-validation loss improvement. For Epitome, we found 5 and 0 to be the most optimal patience and min-delta, respectively, after tuning and comparing the hyper-parameters shown in Supplementary Figure S15(b). Training is halted once either the model begins to converge (based on the min-delta and/or patience hyper-parameters) or the model trains on the maximum number of training steps.

Supplementary Figure S15(a) compares the model accuracy (auPRC) with and without the early stopping method on 85 ChIP-seq targets. Epitome early stopping validation performs comparably with running Epitome for a set number of 5000 training batches, with the exception of ZNF274. ZNF274 achieves significantly better performance than its baseline method because it requires an average of 2,680 batches using early stopping validation, while other ChIP-seq targets require a mean of 4,300 batches using early stopping validation. These results imply that ZNF274 benefits from early stopping more than other ChIP-seq targets, and thus avoids overfitting that occurs when running for 5,000 batches.

#### 4.3.6 Adding histone modifications to the CASV

Figures 4(c) and 4(d) demonstrate the effect on performance of adding histone modifications in addition to DNase-seq to compute cell type similarity in the CASV. We extended Equation 3 to compute the similarity of multiple assays between two cell types. To extend the CASV to use additional histone modifications, we compute the CASV for each assay, as shown in Equation 3, and concatenate results together as input features.

To assess the effect of extending the CASV to include histone modifications, we trained 256 Epitome models on 13 TFs, including: TCF12, MAFK, JUND, YY1, REST, CTCF, GABPA, RFX5, TBP, SIN3A, NRF1, KDM1A, EP300. These TFs were chosen because they had ChIP-seq availability in at least four ENCODE cell lines that also had ChIP-seq for 7 histone modifications. Each model considered a different combination of the following histone modifications to be used in the CASV to compare cell lines: DNase, H3K9ac, H3K4me3, H3K4me2, H3K4me1, H3K36me3, H3K27me3, and H3K27ac. For each of the 256 combinations of histone modifications and DNase-seq considered in the CASV, we trained four models on three of four cell lines and evaluated on the fourth held out cell lines. Cell lines considered include K562, H1-hESC, HepG2, and GM12878. Each model was evaluated on 50,000 regions from chromosome 7.

### 4.4 Comparison to existing methods for predicting transcription factor binding sites in new cellular contexts

We compared Epitome to three methods that are designed to predict TFBS in cellular contexts not seen during training. This comparison includes Avocado [30], Catchitt [24], and DeFCoM [23]. Each of these methods uses a different algorithm and different features as input to train and predict TFBS. Information regarding each algorithm and the features used for each method are listed in Supplementary Table S1. Because each method differs in their features and regions used for training, we compare to each method differently to ensure a fair comparison. We compared these methods by predicting binding sites for 77 TFs and chromatin modifiers in 40 held out cell lines, tissues, and primary cells, enumerated in Supplementary Table S6. To ensure that methods were not tested on sequences that are similar to the training data, we held out chromosomes 8 and 9 for evaluation of all methods. We specifically chose to hold out two whole chromosomes to ensure that there were sufficient chromosomes for training all models, allowing methods to sufficiently generalize. Additionally, this strategy of holding out entire chromosomes for validation and testing was similarly used by competing methods [19, 24] and ENCODE DREAM challenges [36]. However, because Avocado cannot predict epigenetic signal in regions it has not seen during training, we train Avocado on chromosomes 8 and 9. Figure 2 compares these methods against two metrics: area under the precision recall curve (auPRC) and and partial area under the receiver operating characteristic curve (pAUC), using sklearn [53]. We calculate the standardized pAUC [54] over the range of false positive rates ranging from 0 to 0.05 to limit our analysis to regions on the receiver operating characteristic curve with high specificity.

#### 4.4.1 Training, evaluation, and comparison to DeFCoM

DeFCoM [23] is a footprinting method that uses enzymatic cleavage patterns of DNase I from DNase-seq or Tn5 transposase from ATAC-seq to predict TF binding sites. We chose to compare to DeFCoM because it was shown to predict TFBS better than nine other footprinting methods [23]. Unlike Epitome, DeFCoM is limited to using data from a single reference cell type for training, and its feature set is limited to a genomic context of 100bp surrounding a genomic locus of interest. As input, DeFCoM requires the genomic positions of motif sites for all available motifs specific to a TF of interest. To gather positions of motif sites, we used motifs from the Cis-BP database [55]. We additionally used motifs published by Kheradpour et al. [56], available at http://compbio.mit.edu/encode-motifs/ (See Supplementary Table S5 for a list of motifs used). All motifs were processed using FIMO [57] to identify motif sites for each motif of interest. We calculate the FIMO background model as a 0 order Markov model over the hg38 genome.

We next trained DeFCoM models on all combinations of TFs and cell types, listed in Supplementary Table S6, that had available ChIP-seq data and motif data for the combination of interest. For each combination, three hg38 ChIP-seq peak experiments were downloaded from ENCODE and combined to define the set of ChIP-seq peaks. We trained each model on all autosomal chromosomes, except chromosomes 7, 8, and 9, which were held out for validation and testing. We trained on all motif hits for a given TF, as identified using FIMO, where motifs overlapping a ChIP-seq peak were labeled as positive and motifs not overlapping a ChIP-seq site were labeled as negative. We additionally filtered out motifs as suggested by Quach et. al. [23]. This filtering strategy removed any motifs that fell in ENCODE blacklisted regions, available at https://github.com/Boyle-Lab/Blacklist/raw/master/lists/hg38-blacklist.v2.bed.gz, overlapped DNase-seq peaks by less than 10%, or were less than 400bp from chromosome boundaries. We additionally removed inactive motifs that had ChIP-seq fold change over control greater than 1. We furthermore did not consider TF and cell type combinations that had less than 10 active motif sites overlapping ChIP-seq peaks. We downloaded 1 DNase-seq alignment file and 1 DNase-seq peak file for each cell type of interest from ENCODE, filtering out potential DNase-seq experiments that had audit errors and biosample treatments. We installed DeFCoM using PyPI and trained DeFCoM using default parameters. In total, we trained 323 DeFCoM models. auPRC and pAUC scores for both methods can be found in Supplementary Table S7.

DeFCoM does not allow training on more than one cell type. Therefore, DeFCoM can not make predictions in a held out cell type while considering information from multiple available cell types. Therefore, to get predictions in a held out cell type, we used an ensemble method by averaging DeFCoM predictions from all models trained using DNase-seq from the remaining cell types to gather predictions in the held out cell type of interest.

Because DeFCoM is a motif-centric method, it was not designed to predict TFBS in regions not overlapping motif sites. Therefore, we only compare Epitome and DeFCoM in regions overlapping motifs, shown in Figure 2(b) and Supplementary Figure S8. Additionally, DeFCoM trains separate models for each TF of interest. We therefore compare DeFCoM to Epitome by training individual Epitome models for each TF of interest, instead of using a joint Epitome model that predicts binding for multiple TFs.

#### 4.4.2 Defining the test set to compare Epitome, Catchitt, and Avocado

Unlike motif centric methods, Catchitt, Epitome, and Avocado can predict TFBS outside of motif sites. We therefore evaluated these three methods on all 200bp binned regions of chromosomes 8 and 9. However, Epitome can only generate features and make predictions in regions overlapping known epigenetic events. When Epitome predicts in regions not overlapping epigenetic events, it will automatically predict non-binding (0). Therefore, if we include genomic regions outside of this set, which are, by definition, non-binding, Epitome will have artificially high area under the receiver operating characteristic curve, as it will predict these regions as absolute true negatives. In contrast, other methods that can predict in these regions will have increased numbers of false positives, resulting in worse precision than if these regions were not included. This is demonstrated in Supplementary Figure S7, which shows comparisons of pAUC and auPRC when evaluated on all 200bp regions on chromosomes 8 and 9. We therefore only evaluate on 200bp regions of chromosomes 8 and 9 that overlap at least one ChIP-seq peak from the TF being evaluated, or overlap a DNase-seq peak from at least one of the cell types that has measured ChIP-seq for the TF in question. This choice of the testing set has three advantages. The first advantage is that it removes all regions in which Epitome will automatically predict 0 in regions that are truly negative, thus providing an advantage to competing methods. The second benefit of sub-sampling negative regions from the genome in general is to balance out positives and negatives to get more meaningful receiver operating characteristic curves. In the case that we consider all non binding regions, these metrics will be skewed heavily towards predicting negatives, and would thus give us less information on the ability of individual methods to detect true positives. The third benefit is that this sampling strategy of the test set isolates evaluation of methods to regions of the genome that are generally harder to predict in. While it is generally a simple classification problem to predict nonbinding in genomic loci with no measured regulatory activity, it is harder to predict nonbinding in regions that have some epigenetic signal. pAUC and auPRC values used to generate Figure 2 using this sampling technique on chromosomes 8 and 9 can be found in Supplementary Table S8.

#### 4.4.3 Training, evaluation, and comparison to Catchitt

Catchitt is a TF binding prediction method that uses DNase-seq and motifs to predict binding sites. We compared to Catchitt because of its recent success as a co-winner of the DREAM challenge [36]. Although Anchor [51] also performed well in this DREAM challenge, we did not compare to it because Anchor used features that were similar to those used by Catchitt and code for retraining Anchor models was not documented well enough to easily reproduce. We trained Catchitt [24] with a default bin width of 50bp windowed on the hg38 genome. We used all motifs as those used to train DeFCoM, available in Cis-BP database and Kheradpour et al. [55, 56] (Supplementary Table S5). Bigwig DNase-seq reads as well as conservative and relaxed ChIP-seq peak files were downloaded from ENCODE [2]. Catchitt was trained on chromosomes 1-4 with 5 iterations.

Similar to DeFCoM, Catchitt can only train on a single cell type. We therefore use an ensemble technique to predict on a held out cell type by averaging predictions from multiple models trained using DNase-seq data from different cell types. We first trained separate Catchitt models on each cell type and TF combination. To evaluate a new cell type, for a given TF, we averaged predictions from all other models trained on that TF, excluding models trained using DNase-seq from the cell type being evaluated. To get a prediction in a 200bp window, we took the maximum score of all 50bp predictions overlapping the 200bp window in consideration.

Catchitt trains separate models for each TF of interest. We therefore compared Catchitt to Epitome by training individual Epitome models for each TF of interest, instead of using a joint model that predicts multiple TFs.

#### 4.4.4 Training, evaluation, and comparison to Avocado

Avocado [30] is a deep neural network tensor factorization method designed to impute, or fill in, missing epigenetic signal. We compare to Avocado because it was shown to perform better than existing imputation methods [30]. We processed data for Avocado by downloading signal p-value tracks for TF ChIP-seq and read-depth normalized signal for DNase-seq for 40 cell types used for training from ENCODE. All tracks were binned into 25bp windows, which is default window size in Avocado. The signal for each 25bp window was calculated as the arcsinh transformation of the mean signal in each 25bp window.

For each cell type evaluated, we trained a separate model that included all DNase-seq and ChIP-seq information from all other cell types, as well as DNase-seq from the cell type being evaluated. DNase-seq from the evaluation cell type was included to provide cell type specific information from the evaluated cell type. We then applied Avocado to impute missing signal for each held out cell type for the 77 TFs and chromatin modifiers across all 25bp windows of chromosomes 8 and 9. To calculate predictions in a 200bp window, we took the maximum score of all 25bp predictions overlapping the 200bp window in consideration.

Avocado trains a joint model for all TFs of interest. We therefore compared Avocado and Epitome by training 40 joint models for each method, where each model held out ChIP-seq information from a different cell type. Each model was trained using all 77 TFs and chromatin modifiers.

#### 4.4.5 Training, evaluation, and comparison to DeepSEA

We additionally compared Epitome to DeepSEA. DeepSEA cannot predict TFBS in held out cellular contexts that were not seen during training. DeepSEA uses DNA sequence alone to predict TFBS, and thus does not use any information specific to the cellular context being predicted. DeepSEA is a convolutional neural network that predicts TFBS better than existing sequence based support vector machine (SVM) methods [19]. For this reason, we chose to include DeepSEA to represent sequence based methods.

Evaluations of Epitome and DeepSEA are shown in Supplementary Figures S10 and S11. We evaluated these methods on ENCODE data aligned to the hg19 genome, because DeepSEA models trained on hg19 data were available publicly [58]. We evaluated these methods on four held out cell lines available in ENCODE: K562, GM12878, HepG2, and H1-hESC. For each of these cell lines, we evaluated the following 17 TFs: CEBPB, CHD2, EP300, GABPA, JUND, MAFK, MAX, MYC, NRF1, RAD21, REST, RFX5, SRF, TAF1, TBP, and USF2. Because DeepSEA trains a joint model to predict TFBS, we compared to Epitome models trained jointly on all 17 TFs. We evaluated on chromosomes 8 and 9 and trained on the remaining autosomal chromosomes, except chromosome 7, held out for validation. We selected all 200bp regions for evaluation as defined in Methods 4.4.2. pAUC and auPRC values used to generate Supplementary Figure S10 can be found in Supplementary Table S9.

#### 4.4.6 Training and evaluating DeepSEA

We use pre-trained DeepSEA models available in Kipoi [58] to evaluate DeepSEA [19]. For a given genomic region, DeepSEA predicts 919 epigenetic marks from ENCODE, including ChIP-seq and DNase-seq from multiple cell types. Of these 919 predictions, we take predictions for the K562, GM12878, HepG2, and H1-hESC cell lines for the 17 ChIP-seq targets evaluated.

### 4.5 Evaluation of motif overlap of ChIP-seq peaks for 77 transcription factors and chromatin modifiers in 40 cell types

Supplementary Figure S5 shows the ratio of ChIP-seq peaks that do not overlap any motif for a TF or chromatin modifier of interest. We calculated the ratio of ChIP-seq peaks that do not overlap a motif by considering all ChIP-seq peaks for a given TF and cell type combination, windowed in 200bp regions. For each 200bp region with a peak, we tested whether that region overlapped any motifs specific to the TF in question. We considered all motifs specified in Supplementary Table S5. Motifs were processed as described in Methods 4.4.1. We calculated the ratio of ChIP-seq peaks not overlapping a motif by dividing the number of 200bp regions that do not overlap a motif by the total number of 200bp regions containing a ChIP-seq peak. TFs, chromatin modifiers, and cell types evaluated are listed in Supplementary Table S6. Ratios used to generate Supplementary Figure S5 can be found in Supplementary Table S10.

### 4.6 Calculating the effect of adding cell types to Epitome on model performance

Figures 3(a) and 3(c) demonstrate the effect of adding more reference cell types to the Epitome model on auPRC performance. We evaluated the auPRC of 82 ChIP-seq targets in separately trained models. Of the 82 ChIP-seq targets evaluated, 59 included TFs and 23 included histone modifications. For each of these 82 ChIP-seq targets, we trained separate models that contained 2 to *n* cell lines, where *n* is the maximum number of ENCODE cell lines that had available ChIP-seq for a given target. For each cell line count, we run 4 fold cross-validation on 4 separate held out cell lines. The four cell lines evaluated were K562, HepG2, H1, and GM12878. For each held out cell line, a model was trained on the *i* most similar cell lines, where the cell line similarity was computed by the Jaccard similarity between peak-called chromatin accessibility of the two cell lines being compared. We computed the mean auPRC of each of these 4 models, which are visualized in Figures 3(a) and 3(c). Heatmaps shown in Figure 4 were generated using Seaborn [59].

### 4.7 Prediction of H3K27ac in neural differentiation

Figure 5 shows Epitome predictions of H3K27ac peaks using ATAC-seq data in neural differentiation across seven time points. Time points included 0hr, 3hr, 6hr, 12hr, 48hr, and 72hr. ATAC-seq and H3K27Ac ChIP-seq was taken from Inoue et al. (accession number GSE115046) [41]. This data set consisted of seven time points for ATAC-seq and H3K27ac ChIP-seq with 2 replicates per time point. Data was aligned, peak called, and replicates were combined using the ChIP-seq and ATAC-seq ENCODE DCC pipelines. ENCODE optimal ATAC-seq peaks were used in Epitome to predict H3K27ac peaks across the seven time points in 39,000 regions. These 39,000 regions were identified by Inoue et al. [41] as regions that had at least one H3K27ac peak in one of the seven time points [41]. Clusters in Figure 5(a) were taken from [41] and included 2,400 peaks. Values for the heatmap in Figure 5(b) for H3K27ac was calculated based on the normalized read counts overlapping the 2,400 peaks.

#### 4.7.1 ATAC-seq baseline predictor of H3K27ac

Figure 5(a) compares predictions of H3K27ac peaks by Epitome and an ATAC-seq baseline predictor. To construct an ATAC-seq baseline predictor, we used IDR enrichment scores of ATAC-seq peaks to determine whether an H3K27ac peak existed within a given region of the genome. These scores ranged from 0 (indicating no ATAC-seq IDR peak) and 44.40.

#### 4.7.2 Training and Evaluating DeepHistone

DeepHistone is a deep learning method that uses DNA sequence and chromatin accessibility signal to predict seven histone modifications. We leveraged DeepHistone to predict H3K27ac peaks across seven timepoints of neural differentiation, shown in Figure 5(a). We trained DeepHistone using default training data parameters and demonstration data as defined in DeepHistone’s GitHub repository. We next used DeepHistone to predict 39010 H3K27ac peaks using features extended 1000bp surrounding each H3K27ac peak of interest. We used optimal ATAC-seq peak signal processed from ENCODE as chromatin accessibility signal for DeepHistone.

#### 4.7.3 Plotting 0hr and 72hr H3K27ac predictions in neural differentiation

Figure 5(c) shows Epitome predictions at 0hr and 72hr time points in regions that have unique ATAC-seq peaks in each of these time points. 0hr peaks (shown in red) indicate regions with an ATAC-seq peak at 0hrs, but no peak at 72hr. 72hr peaks (shown in black) indicate regions with an ATAC-seq peak at 72hr, but no peak at 0hr. Shared peaks (yellow) indicate Epitome predictions of regions that have ATAC-seq peaks at both 0hr and 72 hr.

### 4.8 Prediction of TFBS using ATAC-seq on DNase-seq trained models

Supplementary Figure S14 shows that Epitome models that are trained on DNase-seq and validated using ATAC-seq perform comparably when validated using DNase-seq. We used ATAC-seq peaks from the A549 and K562 cell lines to validate the performance of Epitome models that were trained using DNase-seq and validated using ATAC-seq. ATAC-seq and DNase-seq peaks for A549 and K562 were downloaded from ENCODE (datasets ENCSR220ASC, ENCSR000E, ENCSR483RKN, and ENCSR000EKP). Datasets are listed in Supplementary Table S4. Because ATAC-seq peaks for both A549 and K562 were aligned to the hg38 genome, we used liftover [60] to lift over peaks to the hg19 genome for input into Epitome. Each model was trained on 17 cell types: Panc1, PC-9, OCI-LY7, MCF-7, Karpas-422, K562, IMR-90, HepG2, HeLa-S3, HEK293T, HCT116, H9, H1, GM23338, GM23248, GM12878, and A549. When testing the performance of A549 and K562, the respective cell line was held out for test. Due to limited availability of ChIP-seq data for validation, we validated the A549 and K562 cell lines on 33 and 128 ChIP-seq targets, respectively. For each cell line, we ran whole genome predictions using both ATAC-seq and DNase-seq. We compared performance using auPRC and pAUC with a 5% FPR cutoff.

### 4.9 Resource Requirements

All experiments were submitted and managed using TORQUE, which submitted jobs to nodes with one of two configurations: (1) 40 Intel(R) Xeon(R) CPU E5-2690 v2 @ 3.00GHz CPI CPUs with 254 RAM, or (2) 48 Intel(R) Xeon(R) Gold 6226 CPU @ 2.70GHz CPUs with 376GB RAM.

Due to resource limitations, we trained all models using CPUs only. However, Epitome is designed to work with a GPU if installed on a machine with a GPU that is compatible with the current TensorFlow version. However, because Epitome spends a majority of its time computing features for input into the model, most of the resources used for Epitome are CPU intensive, and thus a GPU is not required.

## 5 Data Availability

Epitome is a python package that uses many libraries, including: PyRanges [61], Scikit-learn [53], TensorFlow [62]. Libraries Matplotlib [63] and Seaborn [59] are used for figures.

Source code for Epitome can be found at GitHub. Epitome can be installed through pypi. Documentation for Epitome can be found at readthedocs.

All datasets were curated using ENCODE [2, 1] and ChIP-Atlas [35]. All curated datasets for the hg19 and hg38 genomes used to train and validate Epitome models are publicly available on Amazon S3. GEO accession number GSE115046 was used for evaluation of H3K27ac in neural differentiation [41].

## 1 Supplementary Figures

**Figure S1:**
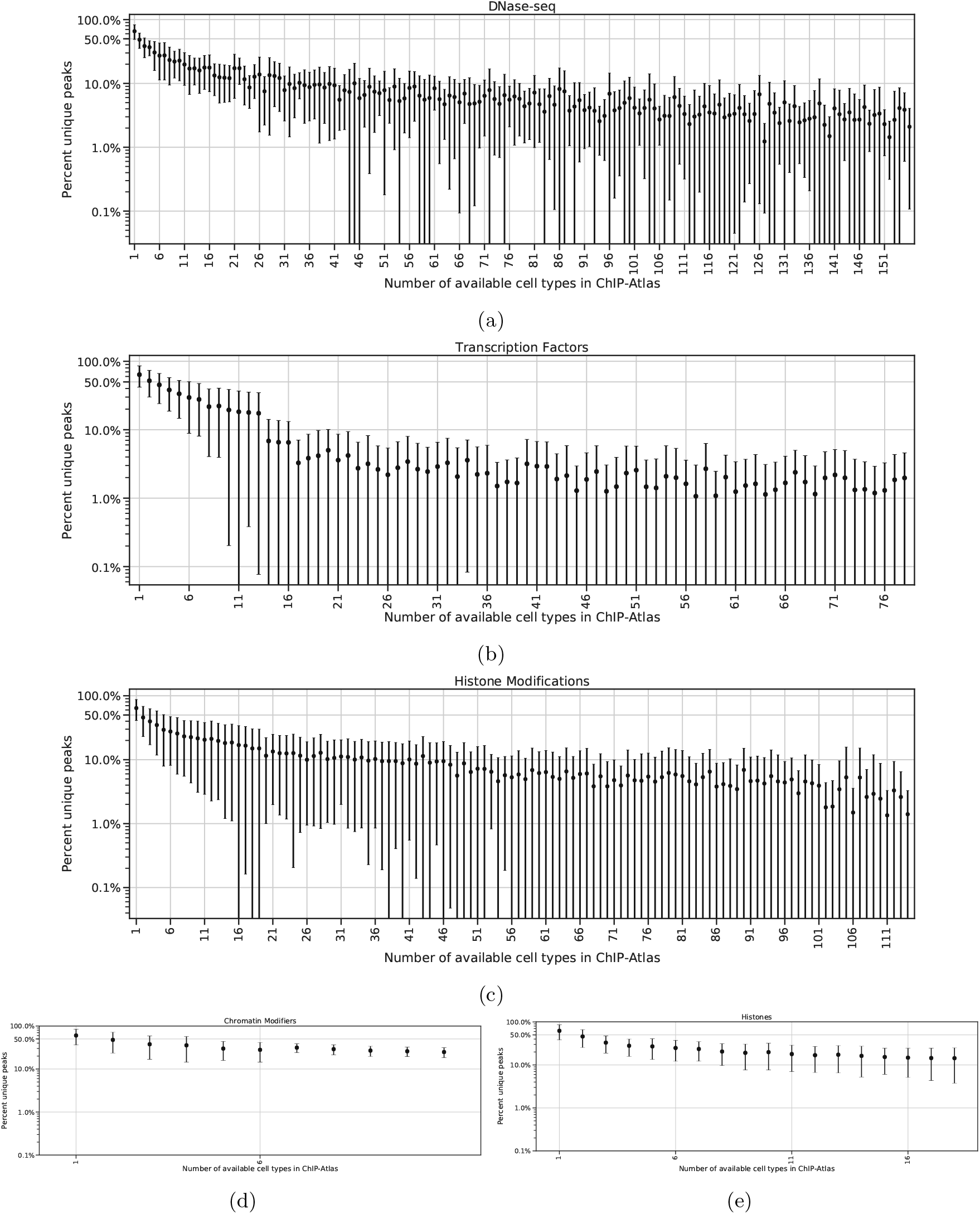
Related to Figure 1(a). Weighted means and standard deviations for percent of unique peaks observed in a cell type as the number of available cell types for a given epigenetic event increases. Means and standard deviations are weighted inversely proportional to the number of datapoints for a given target. Data curated from the ChIP-Atlas database [35]. Results are broken down into the following categories: (a) DNase-seq, (b) transcription factors, (c) histone modifications, (d) chromatin modifiers, and (e) histones.

**Figure S2:**
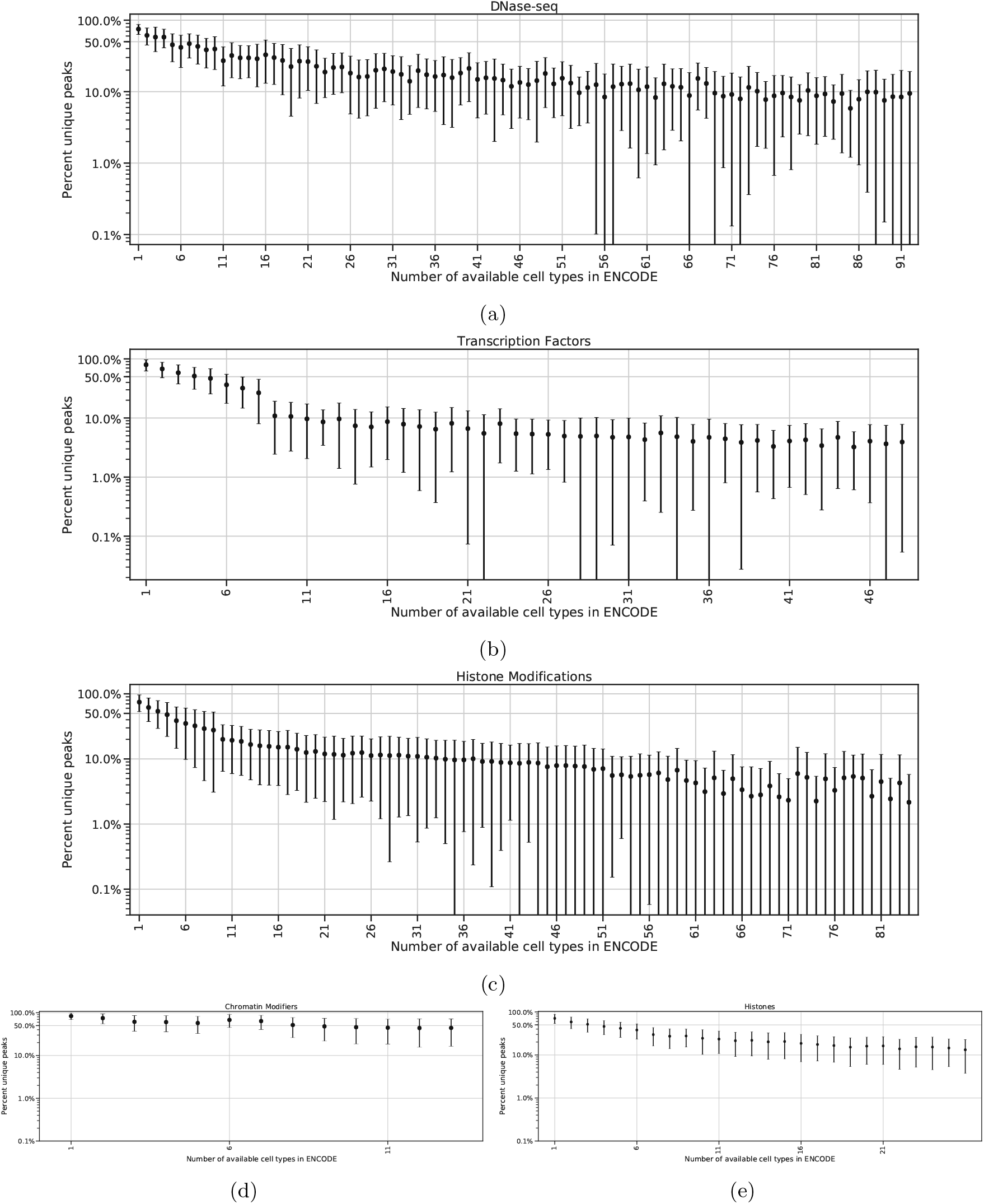
Related to Figure 1(a). Weighted means and standard deviations for percent of unique peaks observed in a cell type as the number of available cell types for a given target increases. Means and standard deviations are weighted inversely proportional to the number of datapoints for a given target. Data curated from ENCODE 3 [1]. Results are broken down into the following categories: (a) DNase-seq, (b) transcription factors, (c) histone modifications, (d) chromatin modifiers, and (e) histones.

**Figure S3:**
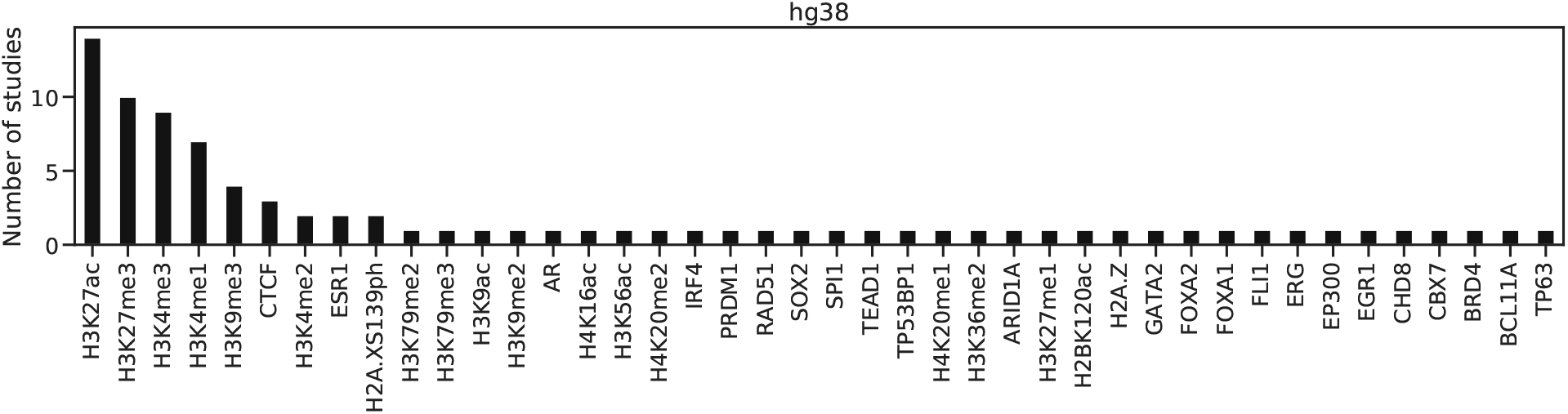
Frequency at which a given ChIP-seq target is present in a study included in ChIP-Atlas [35] that contains at least two ChIP-seq experiments. Only includes studies aligned and processed under the hg38 genome.

**Figure S4:**
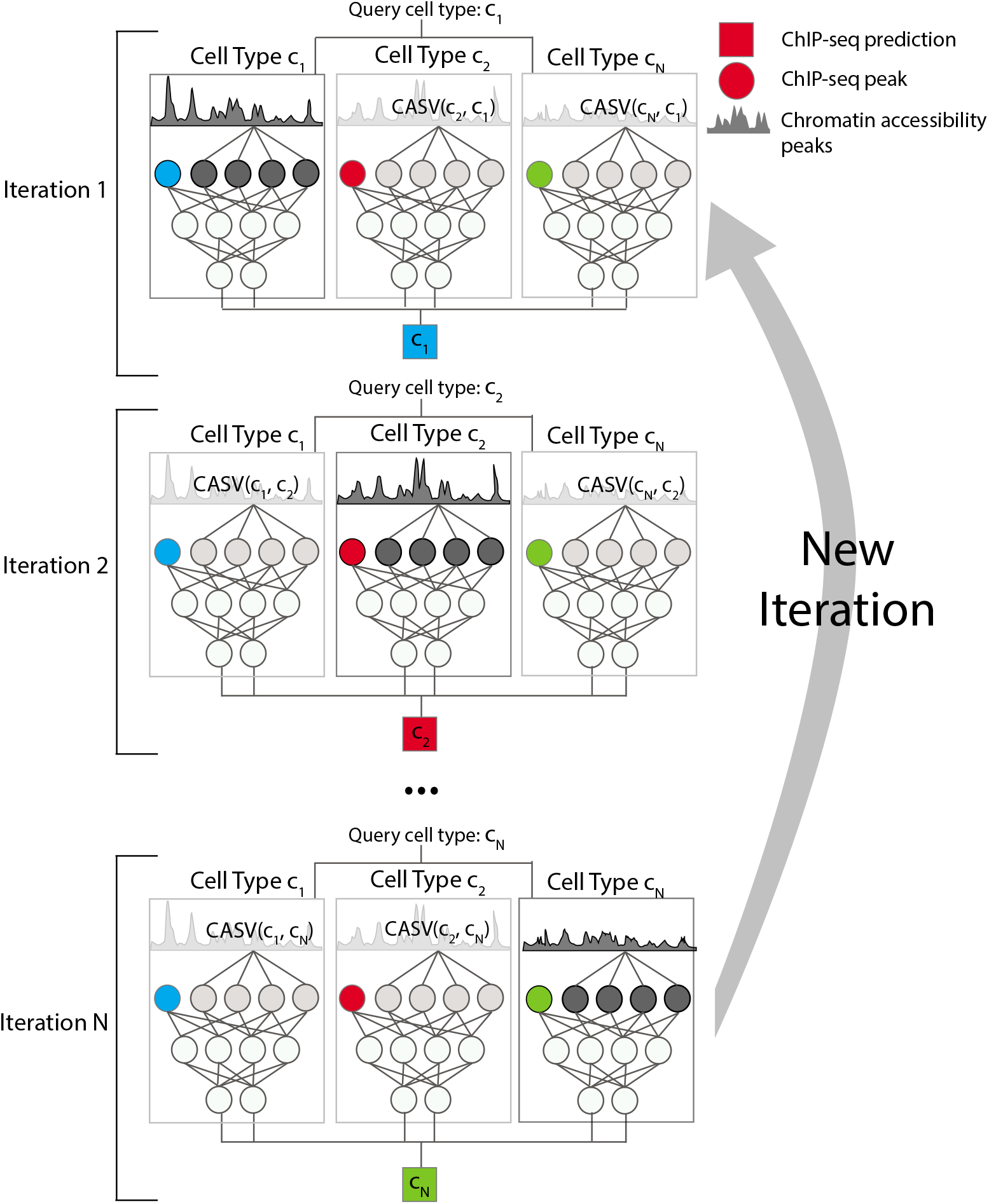
Related to Figure 1. Visual demonstration of cellular setting rotation mechanism used for training Epitome models. Epitome iteratively rotates through which ENCODE cellular setting is used as labels to predict ChIP-seq peaks in a given genomic loci. Remaining cellular settings are used as features. Once all cellular settings in a given genomic loci have been used as labels, a new genomic region is chosen, and all cellular settings are again rotated through to be used as labels.

**Figure S5:**
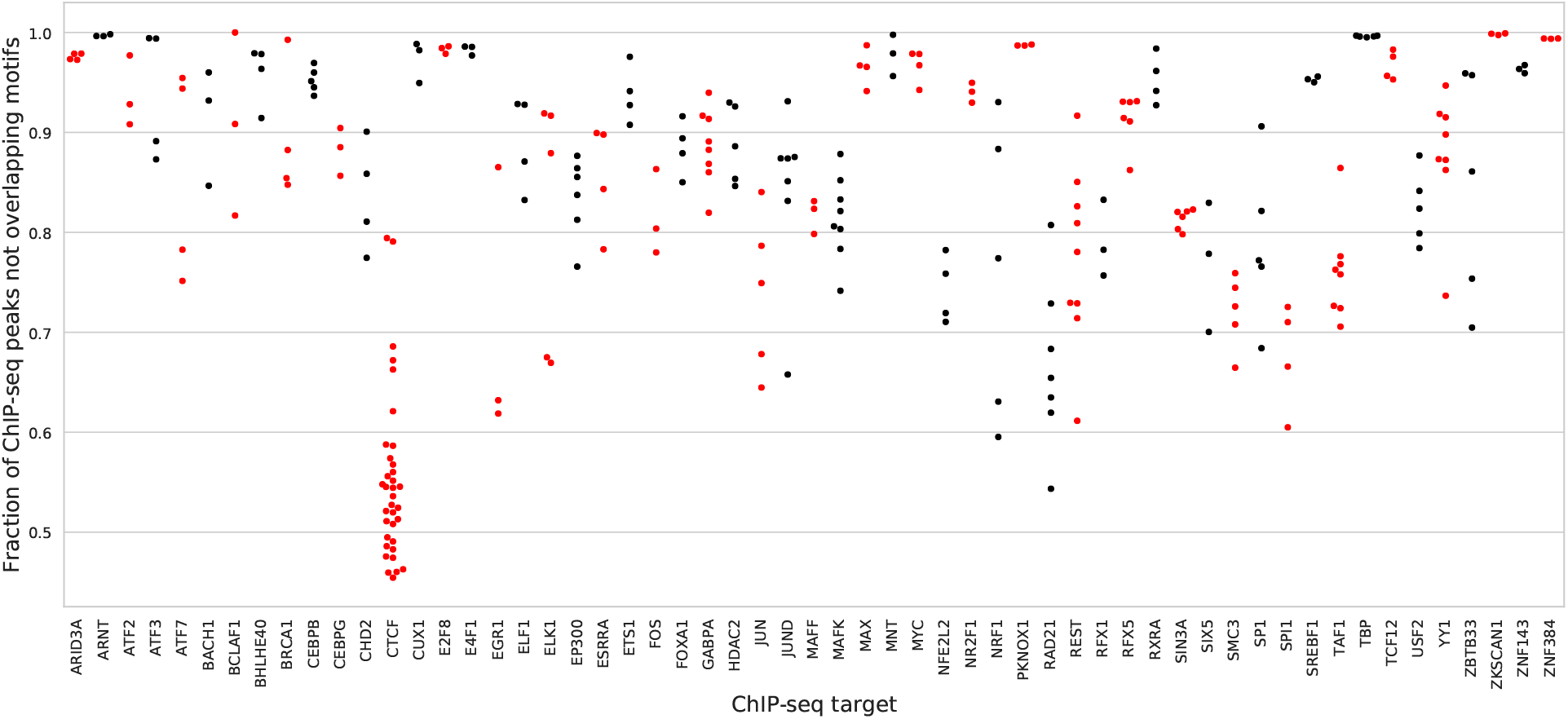
Related to Figure 2. Ratios of ChIP-seq peaks from ENCODE (hg38) that do not overlap any motif. 77 TFs and chromatin modifiers were considered across 40 cell types (Supplementary Table S6). Each data point represents the ratio of motif misses for a ChIP-seq experiment from a TF/chromatin modifier and cell type combination.

**Figure S6:**
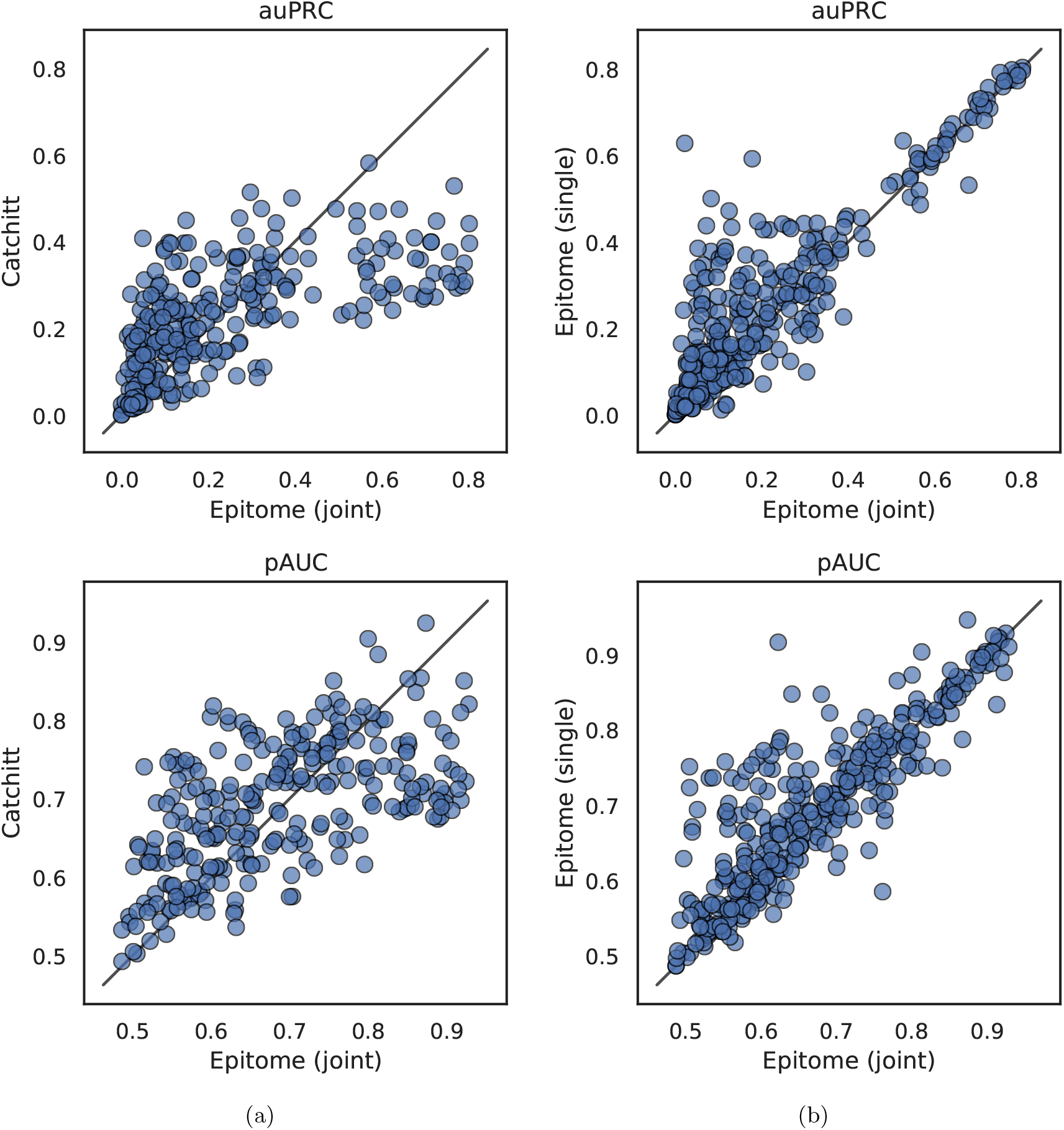
Related to Figure 2. Performance of Epitome joint models, single models, and Catchitt for predicting ChIP-seq peaks for 77 transcription factors on chromosomes 8 and 9 in 40 held out primary cells, tissues, and cell lines. (a) auPRC and pAUC (5% FPR) scores for Epitome joint models and Catchitt. (b) auPRC and pAUC (5% FPR) scores for Epitome models trained jointly and Epitome models trained individually (single) for each TF.

**Figure S7:**
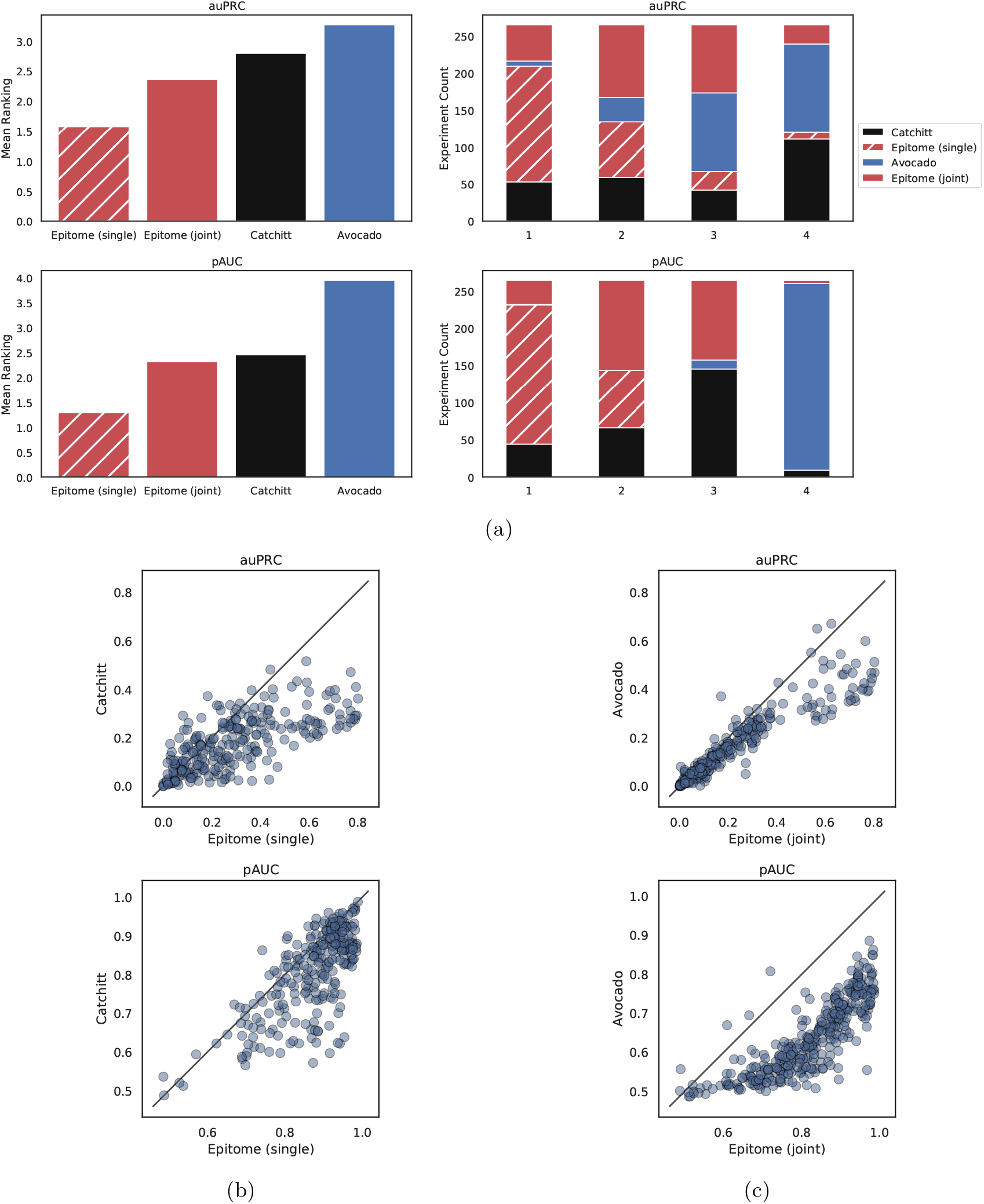
Related to Figure 2. Comparison of Epitome, Avocado, and Catchitt for predicting transcription factor binding sites (TFBS) for 77 transcription factors (TFs) in 40 primary cells, cell lines, and tissues from ENCODE evaluated across all 200bp regions on chromosomes 8 and 9. (a) Frequency at which each method obtains a rank for predicting TFBS across 77 transcription factors in 40 held out cell lines, tissues, and primary cells, totaling 264 comparisons. Evaluated methods include Avocado [30], Catchitt [24], a joint Epitome model, and single Epitome models, where each TF is trained separately. (Left) Mean pAUC (5% FPR) and auPRC ranking for each method. (Right) Frequency at which each method obtains a rank based on pAUC and auPRC. (b) Scatter plots comparing auPRC and pAUC (5% FPR) between Epitome and Catchitt. Both Catchitt and Epitome trained individual models for each TF evaluated. (c) Scatter plots comparing auPRC and pAUC (5% FPR) between Epitome and Avocado. Both Avocado and Epitome trained joint models for all TFs evaluated.

**Figure S8:**
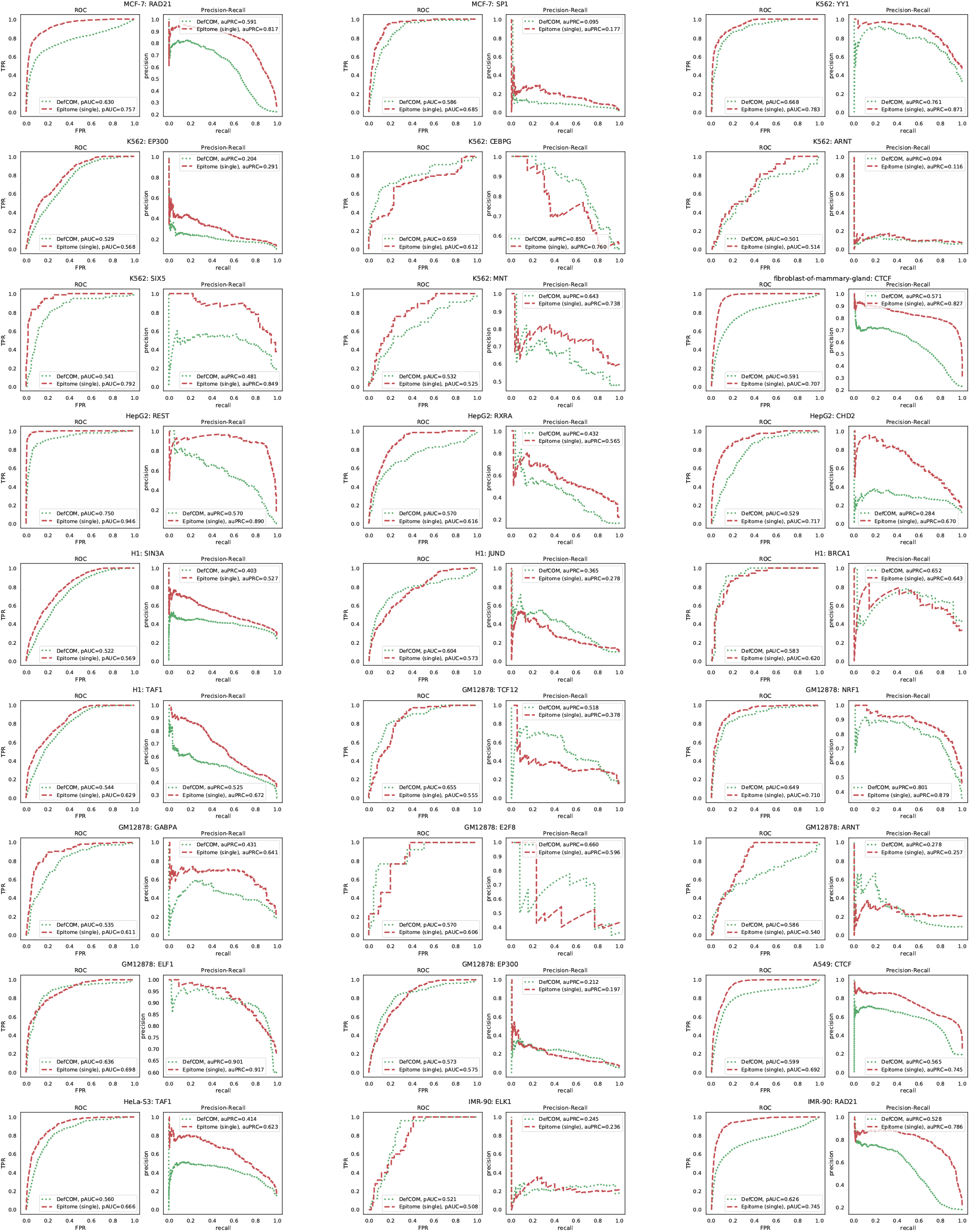
Related to Figure 2. Example ROC and PR curves for Epitome single models and DefCOM. Each plot represents evaluation of a randomly selected transcription factor in a held out cell line, primary cell, or tissue. All models were trained and evaluated on ENCODE processed peaks from the hg38 genome and were evaluated on regions that overlap a motif specific to the TF being evaluated.

**Figure S9:**
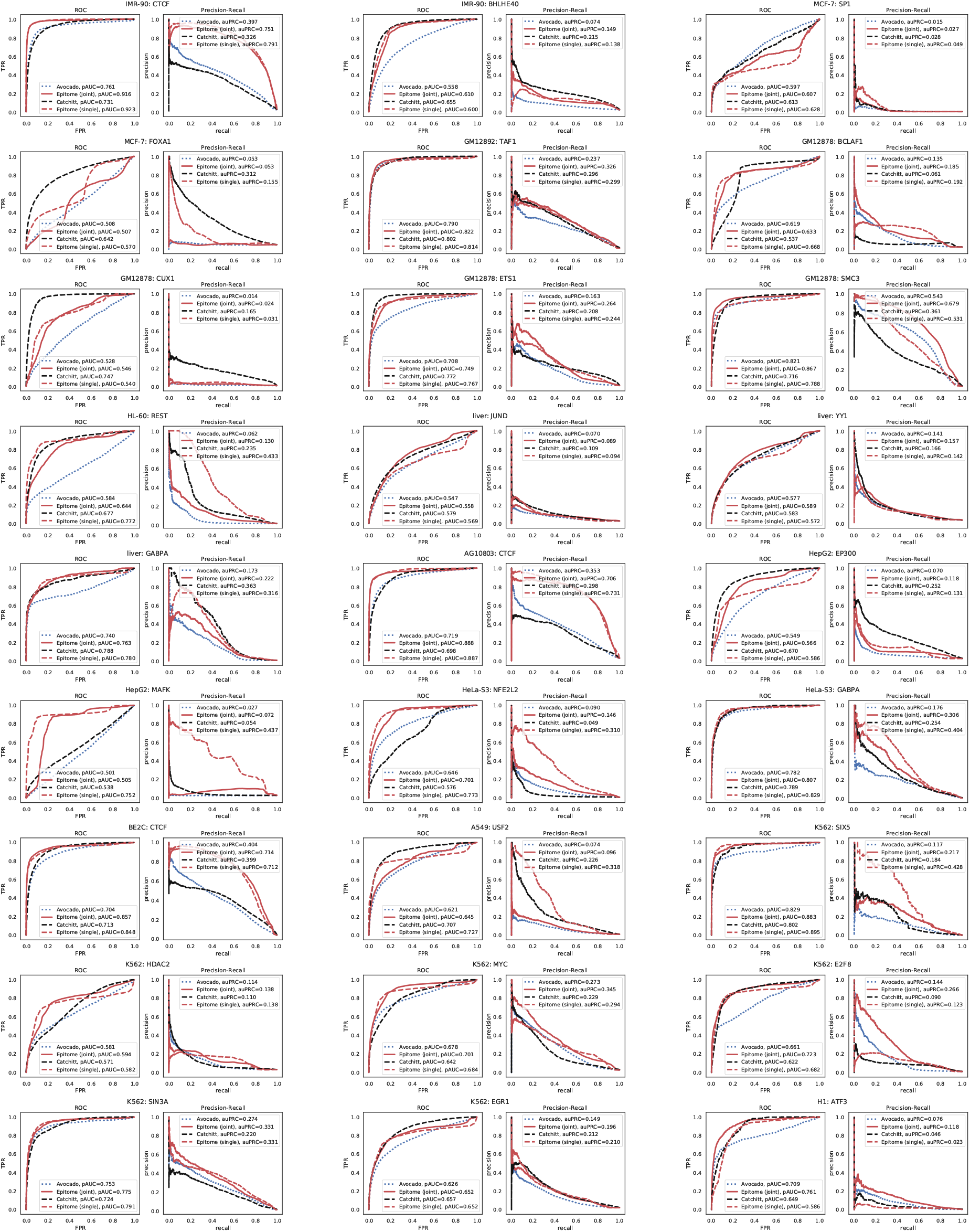
Related to Figure 2. Example ROC and PR curves for Avocado, Catchitt, Epitome single models, and Epitome joint models. Each plot represents evaluation of a randomly selected transcription factor in a held out cell line, primary cell, or tissue. All models were trained and evaluated on ENCODE processed peaks from the hg38 genome and were evaluated on chromosomes 8 and 9.

**Figure S10:**
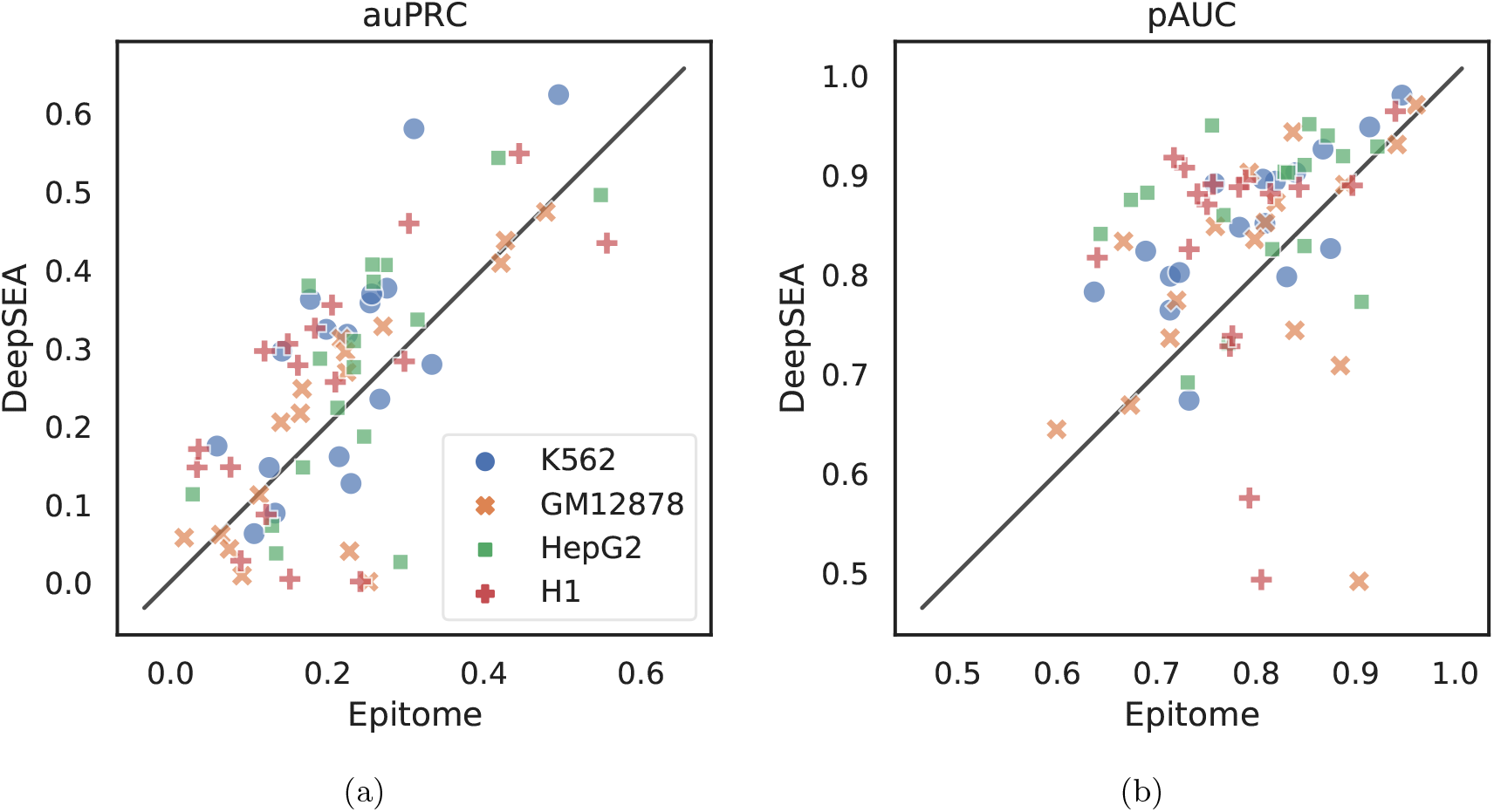
Related to Figure 2. Performance metrics of Epitome and DeepSEA for predicting ChIP-seq peaks for 17 transcription factors on chromosomes 8 and 9 in four held out cell lines, resulting in 68 comparisons. Four held out cell lines include K562, GM12878, HepG2, and H1. Transcription factors compared include: CEBPB, CHD2, CTCF, EP300, GABPA, JUND, MAFK, MAX, MYC, NRF1, RAD21, REST, RFX5, SRF, TAF1, TBP, and USF2. (a) auPRC and (b) pAUC (5% FPR) scores for Epitome and DeepSEA.

**Figure S11:**
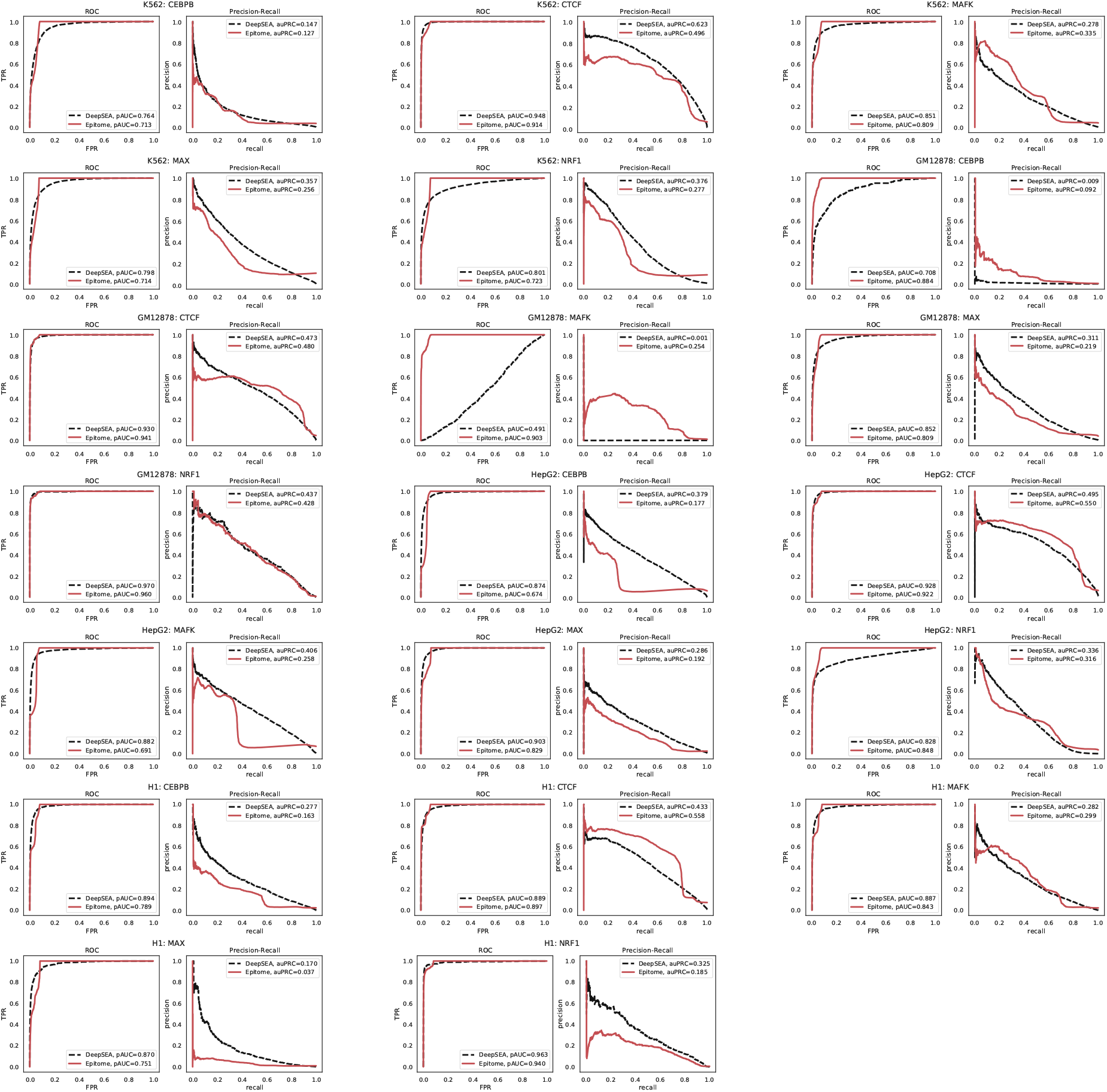
Related to Figure S10. Example ROC and PR curves for DeepSEA and Epitome joint models. Each plot represents evaluation of a randomly selected transcription factor in a held out cell line. All models were trained and evaluated on ENCODE processed peaks from the hg19 genome and were evaluated on chromosomes 8 and 9.

**Figure S12:**
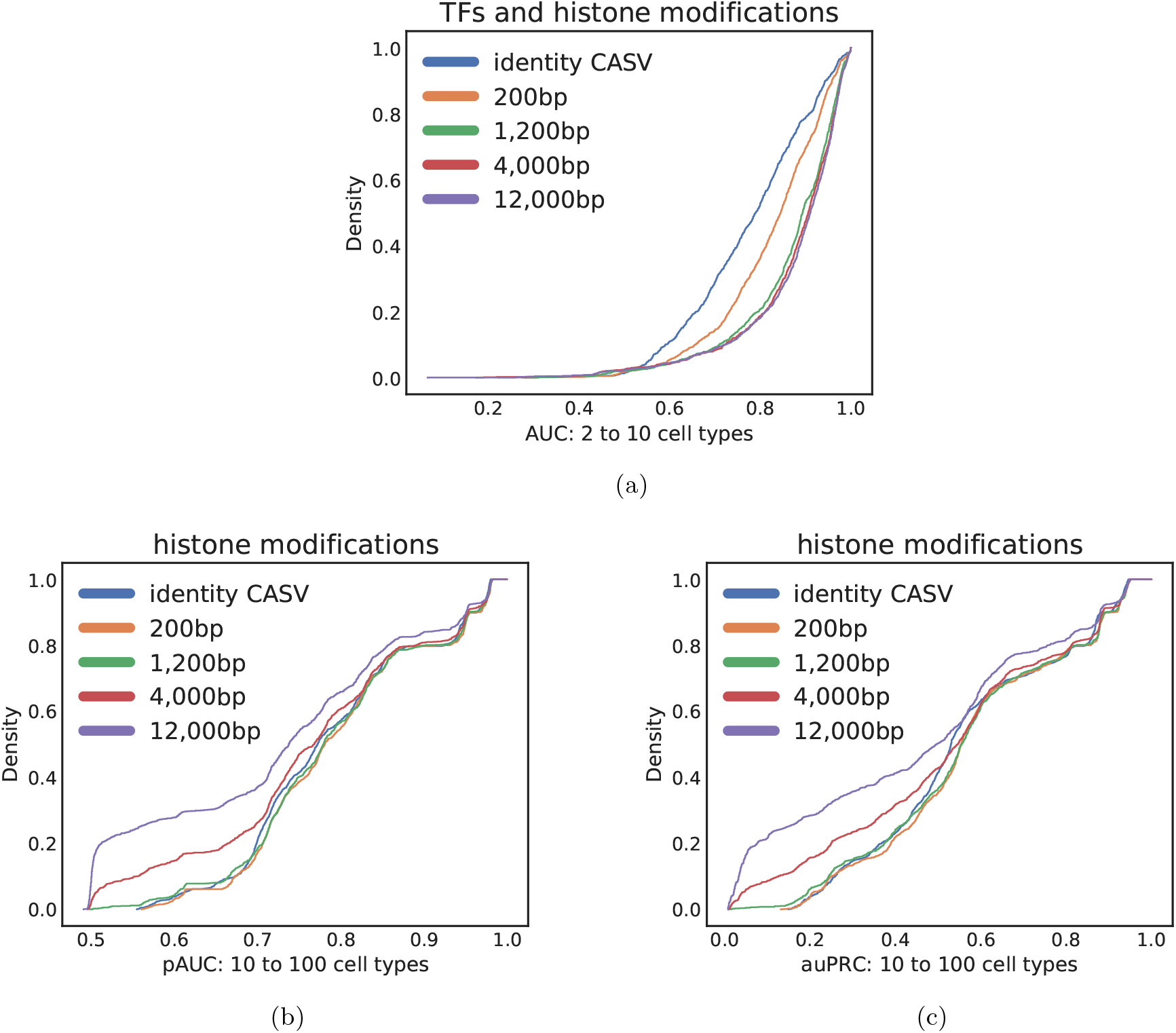
Related to Figure 4. Considering genomic contexts of various sizes to compute cell type similarity in the CASV affects performance of transcription factors (TFs) and histone modifications. Various DNase-seq window sizes are considered for computing the chromatin accessibility vector (CASV). Only DNase-seq is used to compute cell type similarity in the CASV. DNase-seq window sizes considered include no DNase-seq, 200bp, 1,200bp, 4,000bp, and 12,000bp around a peak of interest. (a) Cumulative distribution functions (CDFs) of Epitome performance in terms of area under the receiver operating characteristic curve (AUC) for TFs and histone modifications in Epitome models trained on 2 to 10 cell types. (b),(c) CDFs of Epitome performance for histone modifications in Epitome models trained on more than 10 cell types. Performance was measured in (b) partial area under the receiver operating characteristic curve (pAUC) (5% FPR) and (c) area under the precision recall curve (auPRC).

**Figure S13:**
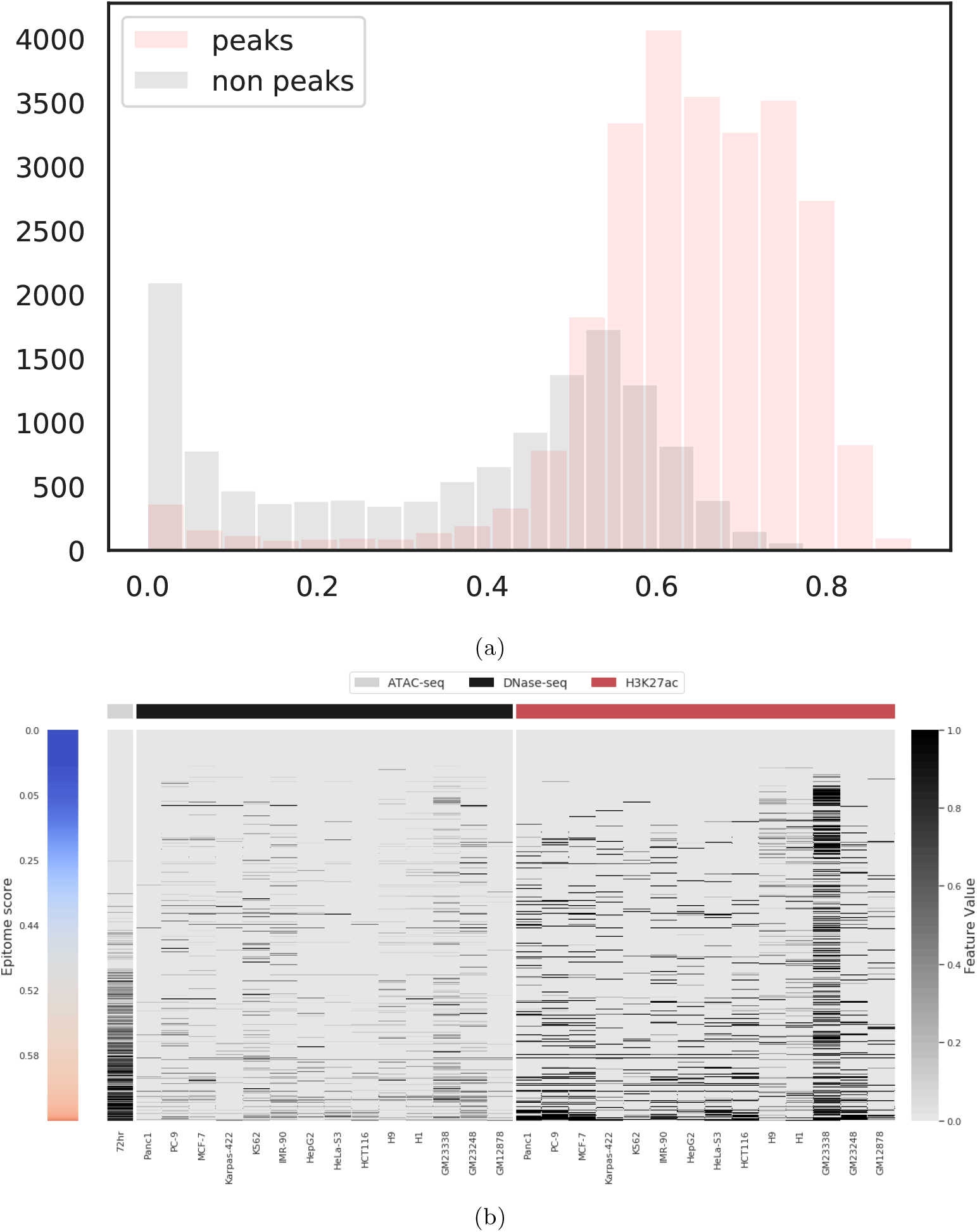
Related to Figure 5. (a) Epitome predictions of H3K27ac peaks at 72hr after neural induction for peak and nonpeak regions. (b) Heatmap of features used by Epitome for 13,248 regions that do not contain H3K27ac peaks at 72hr. Color bar on left represents Epitome scores, where blue represents true negatives and red represents false positives.

**Figure S14:**
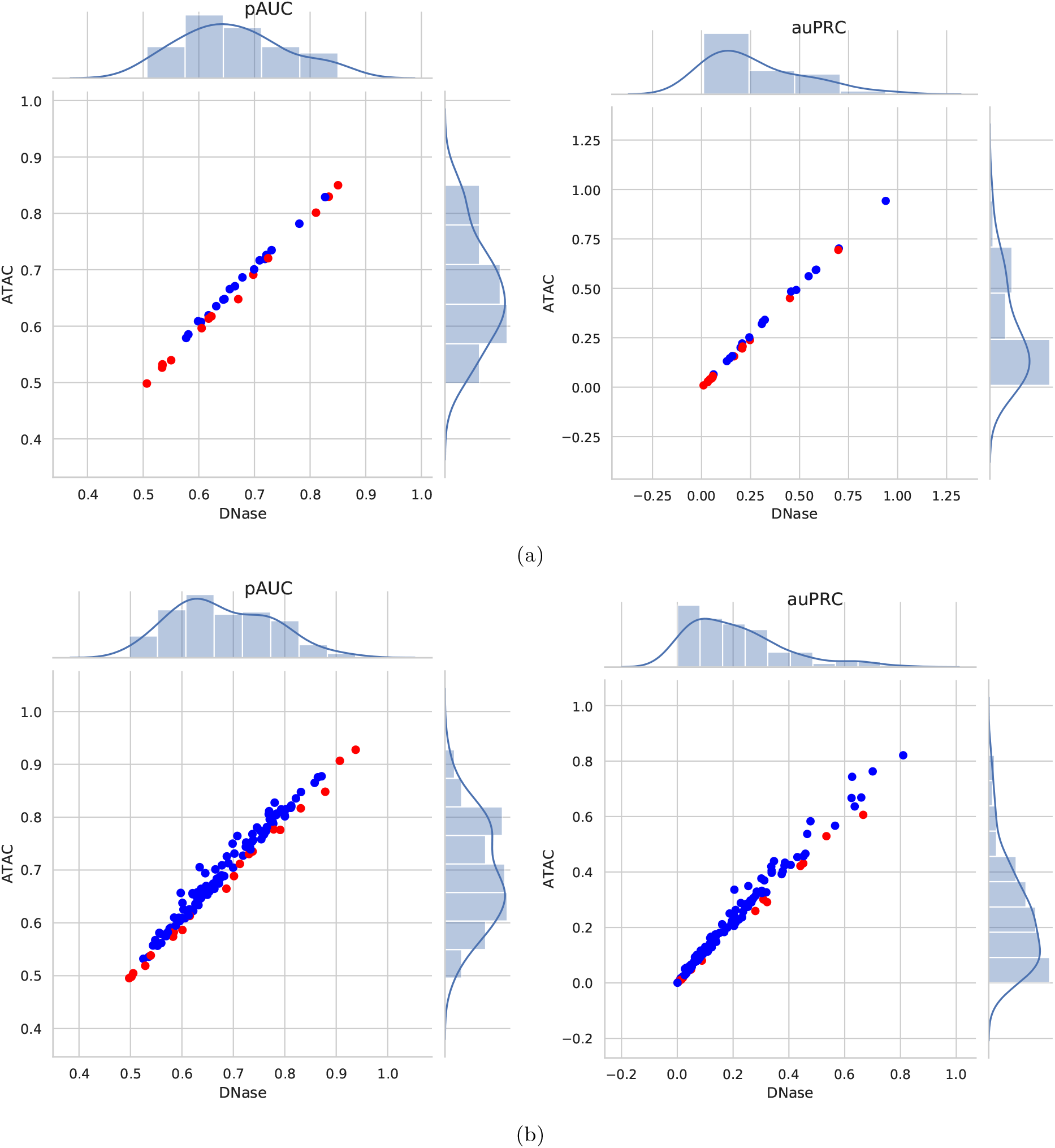
Epitome performs comparably when training models using DNase-seq and evaluating on a new cell line using ATAC-seq. (a) Comparative pAUC (5% FPR) and auPRC performance of 33 TFs when predicting genome wide binding in the A549 cell line using an Epitome model trained using DNase-seq. x axis shows pAUC (left) and auPRC (right) using ENCODE A549 DNase-seq during evaluation, and y axis shows pAUC (left) and auPRC (right) using ENCODE ATAC-seq during evaluation. Blue indicates TFs that perform better when predicted using ATAC-seq data during evaluation. Red indicates TFs that perform better when predicted using DNase-seq data during evaluation. (b) Comparative pAUC (5% FPR) and auPRC performance of 128 TFs when predicting genome wide binding in the K562 cell line using an Epitome model trained using DNase-seq. x axis shows pAUC (left) and auPRC (right) using ENCODE K562 DNase-seq during evaluation, and y axis shows pAUC (left) and auPRC (right) using ENCODE ATAC-seq during evaluation.

**Figure S15:**
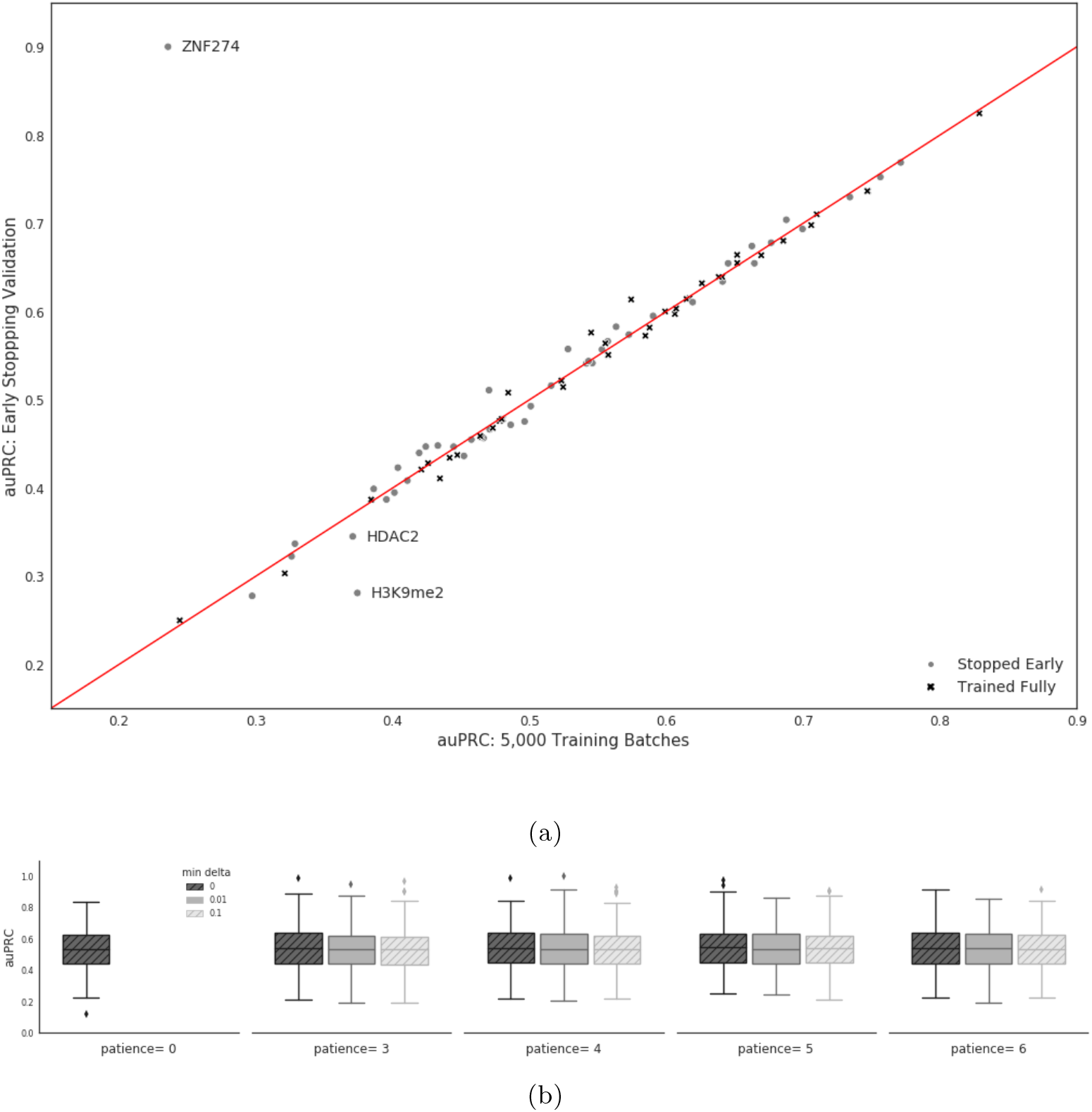
Performance of single TF Epitome models with and without early stopping validation. Values are area under the precision recall curve (auPRC). Models trained without the early stopping validation method trained for 5,000 batches. 5 models were trained for each combination of parameters using early stopping validation method while 5 models were trained without early stop validation. (a) Median auPRC performance of single TF Epitome models with and without early stopping validation, using hyper-parameters of a patience of 5 and minimum delta of 0. The median auPRC is computed across 5 models. 44 out of the 85 TFs stopped training early (before 5,000 batches), with an overall mean of 4,300 training batches. (b) auPRC of single TF Epitome models on different minimum delta and patience hyper-parameters. Each sub-plot indicates a different patience hyper-parameter (the x-axis), and each bar plot hue indicates a different min delta hyperparameter. The first sub-plot with a 0 patience and 0 min delta is the performance from the baseline model, which was trained on 5,000 training batches (without stopping early).

**Figure S16:**
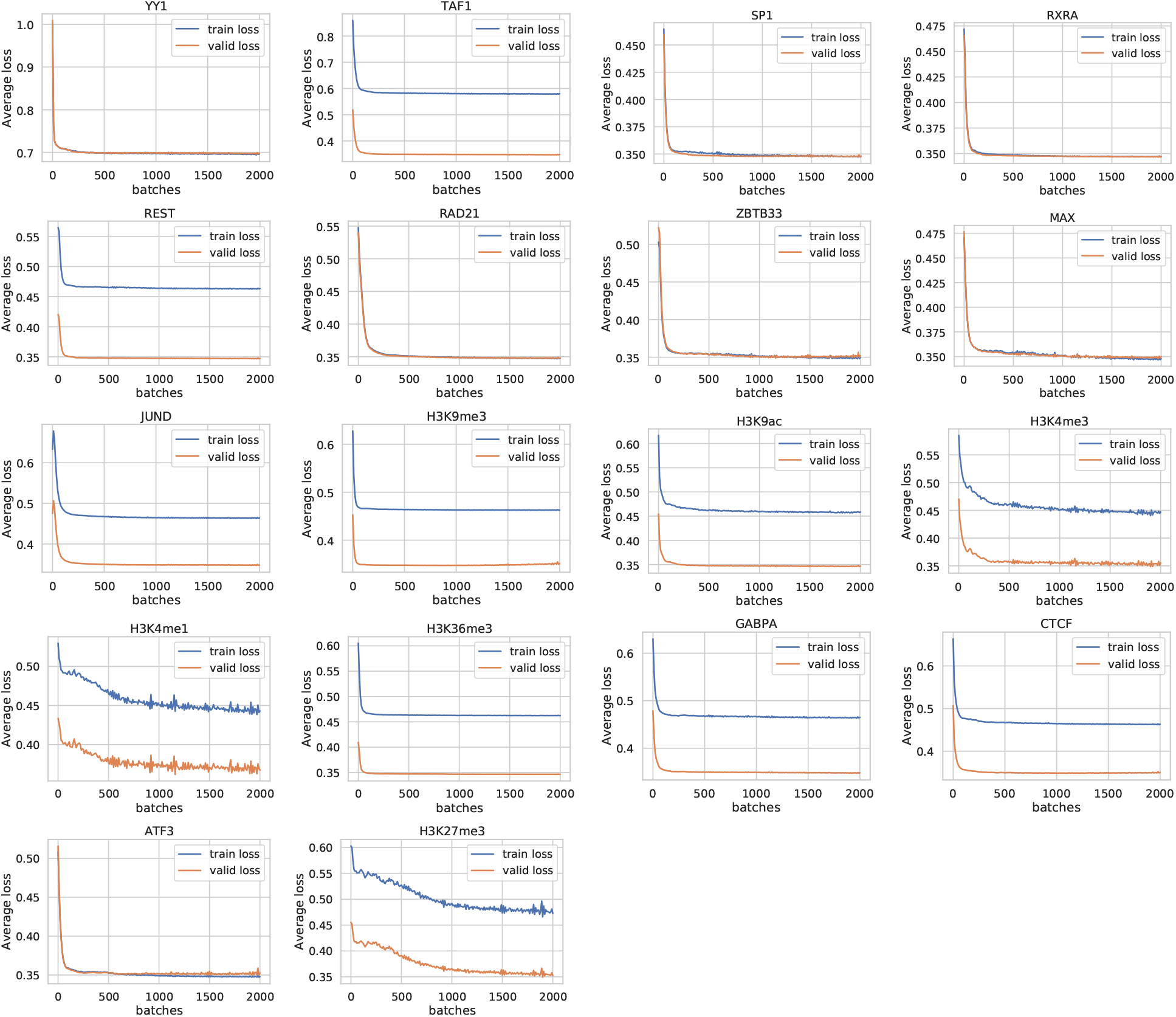
Average training and validation loss for 18 ChIP-seq targets, including transcription factors, chromatin modifiers and histone modifications. All ChIP-seq targets were trained jointly. Average train loss was calculated from 10,000 sampled points from the training dataset. Valid loss was calculated from all points on chromosome 7 meeting the sampling criteria as described in Section 4.3.4. Average loss is calculated as the sigmoid cross entropy, averaged across all evaluated data points (See Equation 2). The model was trained for 2000 iterations, without early stopping.

## 2 Supplementary Tables and Captions

**Table S1:**
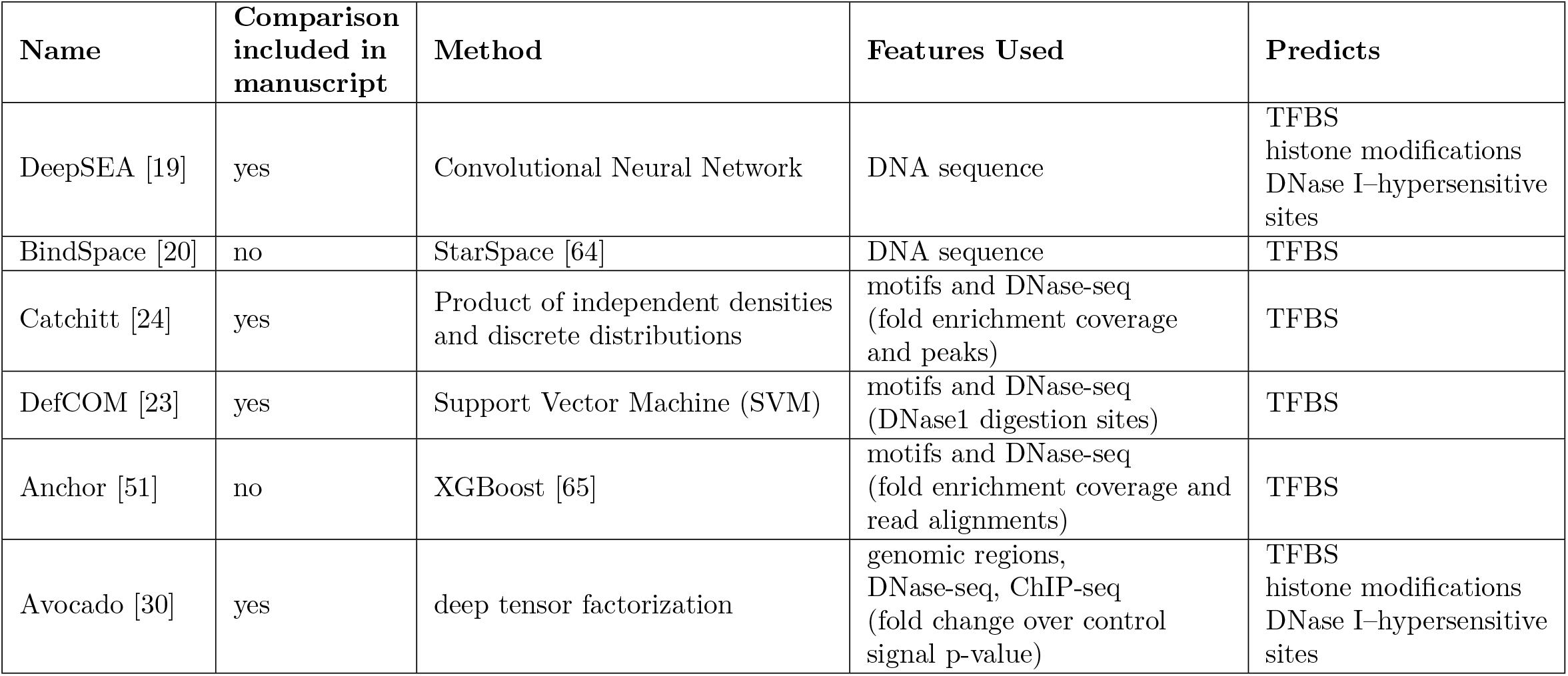
List of methods for predicting transcription factor binding sites (TFBS). Related to Figure 2. Methods compared to include DeepSEA, Catchitt, DefCOM, and Avocado.

| Name | Comparison included in manuscript | Method | Features Used | Predicts |
| --- | --- | --- | --- | --- |
| DeepSEA [19] | yes | Convolutional Neural Network | DNA sequence | TFBS<br>histone modifications<br>DNase I-hypersensitive sites |
| BindSpace [20] | no | StarSpace [64] | DNA sequence | TFBS |
| Catchitt [24] | yes | Product of independent densities and discrete distributions | motifs and DNase-seq (fold enrichment coverage and peaks) | TFBS |
| DefCOM [23] | yes | Support Vector Machine (SVM) | motifs and DNase-seq (DNase1 digestion sites) | TFBS |
| Anchor [51] | no | XGBoost [65] | motifs and DNase-seq (fold enrichment coverage and read alignments) | TFBS |
| Avocado [30] | yes | deep tensor factorization | genomic regions, DNase-seq, ChIP-seq (fold change over control signal p-value) | TFBS<br>histone modifications<br>DNase I-hypersensitive sites |

Table S2: List of ChIP-seq and DNase-seq ENCODE accession numbers used for training and validation of models.

Table S3: List of ChIP-seq and DNase-seq ChIP-Atlas accession numbers used for validation of models and calculating the fraction of unique peaks in Figure 1(a).

Table S4: Accession numbers for ATAC-seq used for validation of models and comparison to DNase-seq trained models. Related to Supplementary Figure S14.

Table S5: Motifs used for training and validating DefCoM [23] and Catchitt [24]. Motifs were taken from Cis-BP [55] and Kheradpour et al. [56].

Table S6: Cell types and ChIP-seq targets used for evaluation of Epitome, Avocado, Catchitt, and DefCoM. Related to Figure 2.

Table S7: Area under the precision recall (auPRC) and partial area under the receiver operating characteristic curve (5% FPR threshold, auROC) for DefCoM and Epitome in regions overlapping motifs.

Table S8: Area under the precision recall (auPRC) and partial area under the receiver operating characteristic curve (5% FPR threshold, auROC) for Epitome, Avocado, and Catchitt. Related to Figure 2.

Table S9:Area under the precision recall (auPRC) and partial area under the receiver operating characteristic curve (5% FPR threshold, auROC) for Epitome and DeepSEA. Related to Figure S10.

Table S10: Number of ChIP-seq peaks that overlap motifs for 77 ChIP-seq targets. Related to Supplementary Figure S5.

